# Mechanism of human PINK1 activation at the TOM complex in a reconstituted system

**DOI:** 10.1101/2023.12.23.573181

**Authors:** Olawale G. Raimi, Hina Ojha, Kenneth Ehses, Verena Dederer, Sven M Lange, Cristian Polo Rivera, Tom D. Deegan, Yinchen Chen, Melanie Wightman, Rachel Toth, Karim P. M. Labib, Sebastian Mathea, Neil Ranson, Rubén Fernández-Busnadiego, Miratul M. K. Muqit

## Abstract

Loss of function mutations in PTEN-induced kinase 1 (PINK1) are a frequent cause of early-onset Parkinson’s disease (PD). Stabilisation of PINK1 at the Translocase of Outer Membrane (TOM) complex of damaged mitochondria is a critical step for its activation. To date the mechanism of how PINK1 is activated in the TOM complex is unclear. Herein we report co-expression of human PINK1 and all seven TOM subunits in *Saccharomyces cerevisiae* is sufficient for PINK1 activation. We use this reconstitution system to systematically assess the role of each TOM subunit towards PINK1 activation. We unambiguously demonstrate that the TOM20 and TOM70 receptor subunits are required for optimal PINK1 activation and map their sites of interaction with PINK1 using AlphaFold structural modelling and mutagenesis. We also demonstrate an essential role of the pore-containing subunit TOM40 and its structurally associated subunits TOM7 and TOM22 for PINK1 activation. These molecular findings will aid in the development of small molecule activators of PINK1 as a therapeutic strategy for PD.

## INTRODUCTION

Autosomal recessive mutations in PTEN-induced kinase 1 (PINK1) and the RING-IBR-RING (RBR) ubiquitin E3 ligase Parkin are causal for early-onset Parkinson’s disease (PD) [1, 2]. Cell-based studies have demonstrated that these proteins function together in a common mitochondrial quality control pathway [3–5]. Active PINK1 phosphorylates both Parkin and ubiquitin at an equivalent Serine65 (Ser65) residue [6–9], resulting in activation of Parkin via a feed-forward mechanism triggering ubiquitin-dependent elimination of damaged mitochondria by autophagy (mitophagy) [3–5]. Active PINK1 also indirectly induces the phosphorylation of a subset of Rab GTPases including Rab 8A at a highly conserved Serine residue (Ser111) that lies within the RabSF3 motif [10, 11]. PINK1 encodes a 581 amino acid Ser/Thr protein kinase containing an N-terminal canonical mitochondrial targeting sequence (MTS) (residues 1-34); catalytic kinase domain containing three unique loop insertions (residues 156-513); and N-terminal and C-terminal extensions (NTE, residues 111-132; CTE residues 514-581) that flank the kinase domain [12–14]. The majority of PD-associated mutations are located within the kinase domain highlighting the protective role of PINK1 kinase activity against the development of PD [15, 16].

Under basal conditions newly translated PINK1 protein is rapidly imported into mitochondria through the Translocase of Outer Membrane (TOM) complex whereupon it undergoes consecutive N-terminal cleavage by matrix MPP proteases and the inner mitochondrial membrane PARL protease followed by retro-translocation into the cytosol and degradation by the 26S proteasome [17–20]. Upon mitochondrial membrane depolarisation that can be induced by mitochondrial uncouplers, for example Antimycin A / Oligomycin, PINK1 import is blocked leading to the accumulation/stabilisation of full-length PINK1 at the outer mitochondrial membrane (OMM) [21–24] and catalytic activation [8, 25]. Mitochondrial depolarisation promotes PINK1 stabilisation at the TOM complex that can be visualised by blue native PAGE (BN-PAGE) as a ∼700 kDa complex distinct from the native TOM complex that migrates as a ∼500 kDa band [26, 27].

The TOM complex is composed of seven subunits, TOM 5, 6, 7, 20, 22, 40 and 70 and is highly conserved through evolution [28–30]. It has been best characterised in yeast for its role in the recognition of mitochondrial precursors synthesised in the cytosol, and their directed import through the pore-containing subunit TOM40 that spans the OMM [31, 32]. Cryo-EM studies of yeast and the human TOM complex have revealed that they form dimeric structures requiring TOM22 binding to TOM40 to form the core complex [33–35]. TOM70 and TOM20 are both loosely associated with the complex and have not been visualised in high resolution cryo-EM structures to date [33–35] although a low resolution 6.8 Å cryo-EM structure of the *Neurospora crassa* TOM complex has been reported with TOM20 [36]. Both TOM70 and TOM20 contain large hydrophilic domains that extend into the cytosol and act as receptors with overlapping preference for precursor proteins [31, 32]. The role of the smaller TOM subunits 5, 6, and 7 have been shown in yeast to be important for assembly of the core complex but their role in mammalian cells has been less well understood [30–32, 37].

Despite considerable research in the field, the mechanism of mammalian PINK1 activation at the TOM complex remains incompletely understood [3]. In a genetic screen, it was discovered that the TOM7 subunit played an essential role for PINK1 stabilisation and activation at the OMM [38, 39]. We and others recently found an intramolecular interaction between the PINK1 NTE and CTE regions [12–14] and that this is required for stabilisation of human PINK1 to the TOM complex [13, 14] and its subsequent activation by autophosphorylation at Ser228 [13]. Numerous pathogenic PD mutations are located within the NTE:CTE interface that prevent human PINK1 recruitment to the TOM complex highlighting the importance of this activation mechanism to disease [13, 14].

Due to the essential role of the TOM complex in mammalian cells, a systematic analysis of the contribution of TOM subunits for PINK1 activation has not been possible to date and furthermore, it has not been established whether the stabilisation of PINK1 to the TOM complex is necessary or sufficient for its activation. To dissect the molecular mechanisms of PINK1 stabilisation at the TOM complex, we have employed a reconstitution system in which genes encoding human PINK1 and the seven subunits of the human TOM complex have been introduced into the budding yeast, *Saccharomyces cerevisiae,* which do not express PINK1. Strikingly we observe that co-expression of wild-type human PINK1 and the seven TOM subunits is sufficient to reconstitute PINK1 activation and we have exploited this to assess the role of individual TOM subunits towards PINK1 activation. Combining the yeast reconstitution system with AlphaFold modelling and mutagenesis studies in mammalian cells, we propose the mechanism of how PINK1 is stabilised and activated at the TOM complex via interaction with the TOM20 and TOM70 subunits. This systematic analysis provides new fundamental insights into the regulation of human PINK1 that will aid in the development of small molecule activators of PINK1.

## RESULTS

### Co-expression of PINK1 with TOM complex subunits is sufficient for its activation

We initially generated stable transformant strains of the budding yeast, *Saccharomyces cerevisiae*, expressing human wild-type or kinase-inactive (D384A) full length PINK1-3FLAG with or without all seven subunits of the human TOM complex (TOM70, residues 1-608; TOM40, residues 1-361; TOM22, residues 1-142; TOM20, residues 1-144; TOM7, residues 1-55; TOM6, residues 1-64; TOM5, residues 1-51) (Fig. 1A and fig. S1A-B). Following induction of protein expression with 2% galactose, PINK1 pathway activity was determined by immunoblotting of whole-cell yeast extracts for endogenous ubiquitin phosphorylation or human PINK1 Ser228 (trans)autophosphorylation. In the yeast strain expressing human PINK1 alone we did not observe significant activation of PINK1 (Fig. 1B-C) consistent with previous studies showing that recombinant human PINK1 expressed in *E. coli* or insect cells displays little catalytic activity [40] and that transiently transfected PINK1 alone in human cell lines exhibits low levels of activity under basal conditions [41]. Strikingly we observed that induced co-expression of wild-type human PINK1 together with all TOM complex subunits, led to robust increase in ubiquitin phosphorylation (pS65 Ub) and PINK1 Ser228 autophosphorylation (pS228 PINK1) associated with a significant increase in the levels of PINK1 protein compared to expression of PINK1 alone (Fig. 1B-C). This was not observed when kinase-inactive PINK1 was co-expressed with all TOM subunits (Fig. 1B-C). We further confirmed this in 4 independent yeast strains expressing wild-type PINK1 with or without TOM complex components (fig. S2A-B). This data suggests that PINK1 stabilisation at the TOM complex is sufficient for PINK1 activation. To validate the relevance of PINK1 activation in this reconstitution system, we next generated stable transformant yeast strains of PD-associated pathogenic mutations of PINK1 that have been previously characterised in mammalian cell-based assays following conditions of mitochondrial damage-induced membrane depolarisation [13] including an NTE region mutant Q126P; ATP binding mutant E240K; Insertion 3 (Ins3) substrate binding mutant G309D; and a CTE region mutant 534_535InsQ (fig. S3A). Consistent with previous studies in mammalian cells [13], we observed complete loss of phosphorylated ubiquitin in all mutants (fig. S3A). Further we observed loss of Ser228 autophosphorylation except for the G309D mutant consistent with the impact of this mutant on substrate binding (fig. S3A). For all strains expressing wild-type or mutant PINK1-3FLAG, we observed two bands of PINK1 corresponding to full-length and N-terminal cleaved protein suggesting that recombinant PINK1 in this system is being targeted to mitochondria (Fig. 1B; fig. S2 & 3A). To confirm this, we performed live cell imaging studies of PINK1-GFP and this demonstrated co-localisation of PINK1 with the mitochondrial marker red CMXRos (Fig. 1D). Immunoblotting analysis of mitochondrial fractions of yeast strains expressing PINK1-3FLAG and the TOM complex also revealed that PINK1 and all human TOM subunits tested were localised in the mitochondrial fraction (fig. S3B).

**Figure 1.**
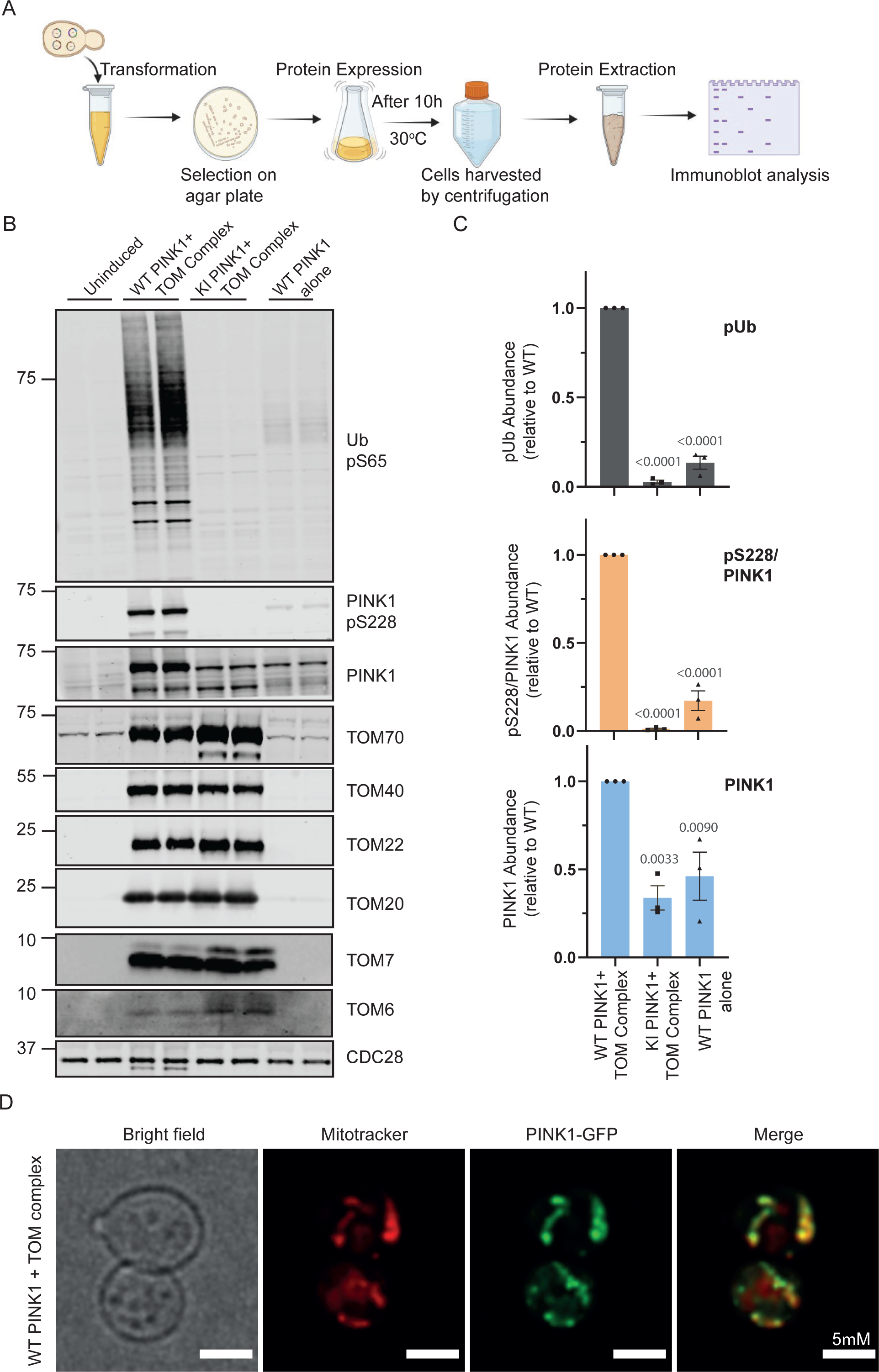
Co-expression of human PINK1 and TOM complex in yeast is sufficient for PINK1 activation. **(A)** Schematic of experimental workflow of PINK1 reconstitution in yeast. Image was created using BioRender.com. Yeast cells were transformed with all four plasmids carrying two plasmids each of the eight components for reconstitution. Cells were selected on a synthetic complete dropout plate, and positive clones were used for protein expression. After expression cells were harvested, lysed and the cell lysate analysed for protein expression. **(B)** Co-expression of human PINK1 and TOM complex subunits induces PINK1 activation. Stable yeast transformants were selected expressing wild-type (WT) or kinase-inactive (KI, D384A) full length-human PINK1-3FLAG and TOM 5, 6, 7, 20, 22, 40, 70 subunits (TOM Complex) or WT human PINK1 alone. Expression was induced by supplementing the growth medium with 2% galactose. 20 µg of whole cell lysates was run on 4-12% Bis-Tris gel and transferred onto nitrocellulose membrane followed by immunoblotting with anti-Ub pS65, anti-PINK1 pS228, anti-total PINK1, and other indicated antibodies. Data representative of 3 independent experiments. **(C)** Quantification of the levels of Ub pS65, PINK1 pS228/PINK1 and total PINK1. Data represents three independent experiments. Statistical analysis was done by ordinary one-way ANOVA where p values relative to WT are shown above the bars. **(D)** Localisation of expressed human PINK1 to yeast mitochondria. Stable yeast transformants expressing wild-type (WT) full length-human PINK1-GFP and TOM 5, 6, 7, 20, 22, 40, 70 subunits (TOM Complex) were generated and mitochondria stained by addition of 500 nM of MitoTracker CMXRos Red. Following incubation, cells were briefly spun down, washed twice with PBS and applied to Concavalin A-coated coverslips which were placed on glass-slides for immediate image acquisition using a Leica DMi8 microscope. Further processing was carried out in the Leica LAS X software platform which includes histogram adjustment and denoising with THUNDER. Images correspond to brightfield microscopy, mitochondria stained by Mitotracker (red) and PINK1-GFP (green).

### Genetic determination of TOM subunits regulating PINK1 activation

We next undertook systematic genetic analysis of the role of the TOM complex on PINK1 activation by generating stable transformant strains missing each of the seven subunits of the TOM complex co-expressed with wild-type human full length PINK1-3FLAG (Fig. 2A). Immunoblotting analysis of PINK1 activation for all strains revealed that the removal of TOM5 or TOM6 subunits did not significantly impair PINK1 activation as assessed by ubiquitin phosphorylation and PINK1 Ser228 autophosphorylation (Fig. 2A-B; fig, S4-S5). Consistent with the previous reports of Youle [38, 39], we observed that removal of TOM7 largely abolished PINK1 activation, and furthermore, had a similar impact to removal of TOM40 or TOM22, and these findings were confirmed in four independent yeast strains for each genotype (Fig. 2A-B; fig, S4-S5). The similarity of the defect of TOM7 to TOM22 is consistent with their critical roles in the assembly and maintenance of the core TOM40 pore-containing complex [33–35]. Interestingly we also observed a reduction in PINK1 activation following removal of the accessory receptor subunits TOM20 or TOM70 and the activation was further reduced in strains lacking both TOM20 and TOM70 (Fig. 2A-B; fig, S4-S5). Overall, our findings indicate TOM subunits can be stratified into three groups based on their effect on PINK1 activation in the reconstitution system: Group 1 (no effect): TOM 5, 6; Group 2 (essential for core complex assembly): TOM 7, 22, 40; and Group 3 (regulatory receptor binding role): TOM 20, 70.

**Figure 2.**
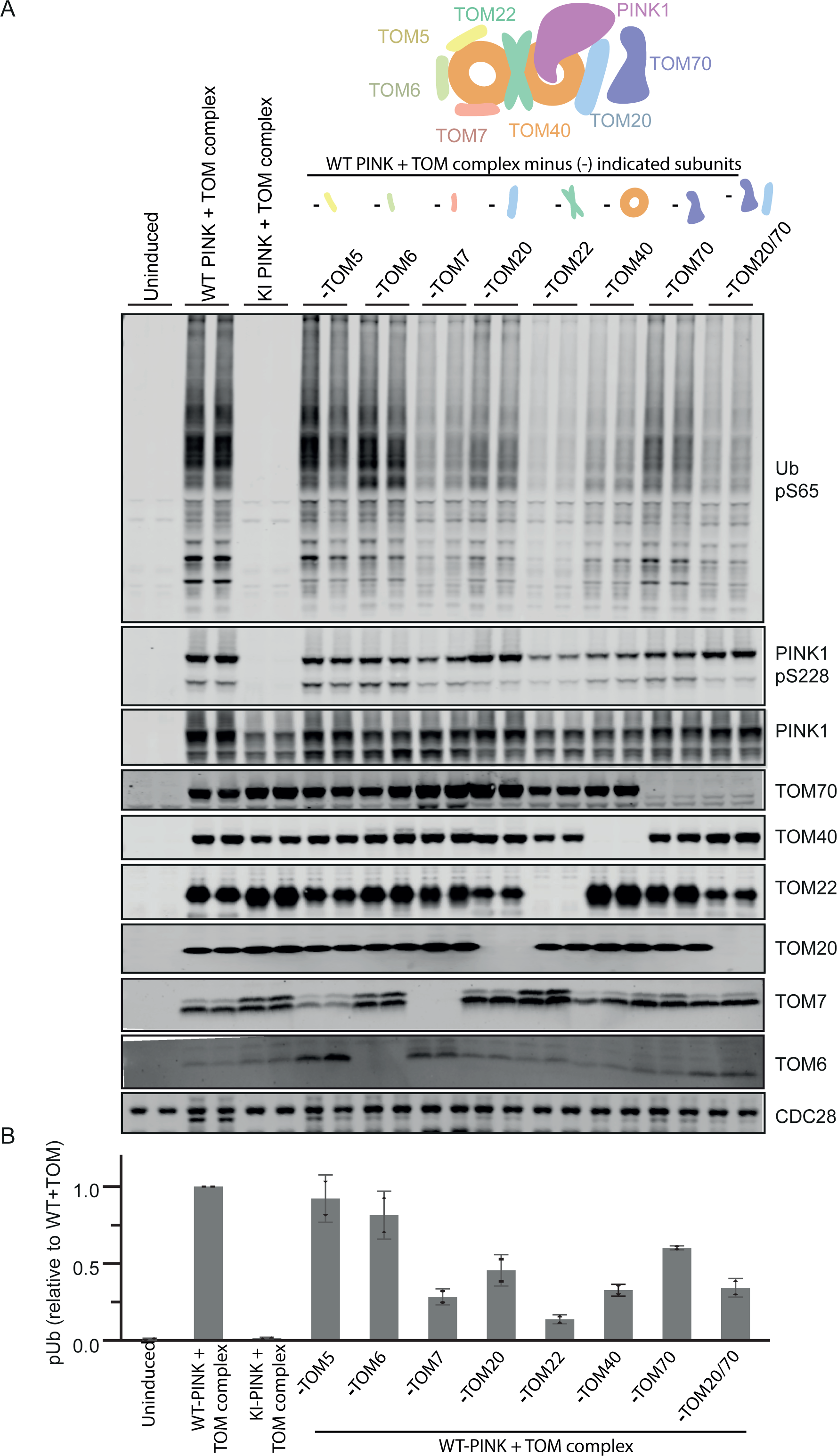
Intact TOM complex required for optimal PINK1 activation. **(A)** Genetic analysis of role of TOM subunits on PINK1 activation. Stable yeast transformants were selected expressing wild-type (WT) or kinase-inactive (KI, D384A) full length-human PINK1-3FLAG and TOM 5, 6, 7, 20, 22, 40, 70 subunits (TOM Complex or TOM complex minus indicated subunits). Expression was induced by supplementing the growth medium with 2% galactose. 20 µg of whole cell lysates was run on 4-12% Bis-Tris gel and transferred onto nitrocellulose membrane followed by immunoblotting with anti-Ub pS65, anti-PINK1 pS228, anti-total PINK1, and other indicated antibodies. Data representative of 2 independent experiments. **(B)** Quantification of Ub pS65 levels normalized to PINK1 expressed with intact TOM complex subunits. Data represent mean+SEM of two independent experiments.

### Structural modelling predicts NTE:CTE interface of PINK1 binds TOM20

TOM20 and TOM70 are not visible on high resolution cryo-EM structures of the yeast and human TOM complex indicating that these subunits are likely to be highly dynamic within the complex [33–35]. To investigate how PINK1 stabilisation at the TOM complex is mediated by TOM20 and TOM70, we employed a locally installed ColabFold notebook [42] to run AlphaFold [43] structure predictions of the PINK1-TOM complex, imputing full-length human PINK1 and varying combinations of TOM subunit sequences in an iterative manner. AMBER structure relaxation [44] was used to ensure appropriate orientation of the side chains and to avoid steric clashes. We were unable to generate a high confidence model of PINK1 and all subunits of the TOM complex together, however, five high confidence models of a single molecule of PINK1, in complex with a TOM dimer containing TOM7, 20, 22, and 40 subunits, were generated based on inter-chain predicted alignment error (inter-PAE) (Fig. 3A-C, fig. S6A-C, Supplementary video). All models correctly predicted the structural interfaces of the core of the TOM complex formed by TOM40, TOM22 and TOM7 in line with existing cryo-EM structures (Fig. 3A-B, fig. S6A-B).

**Figure 3.**
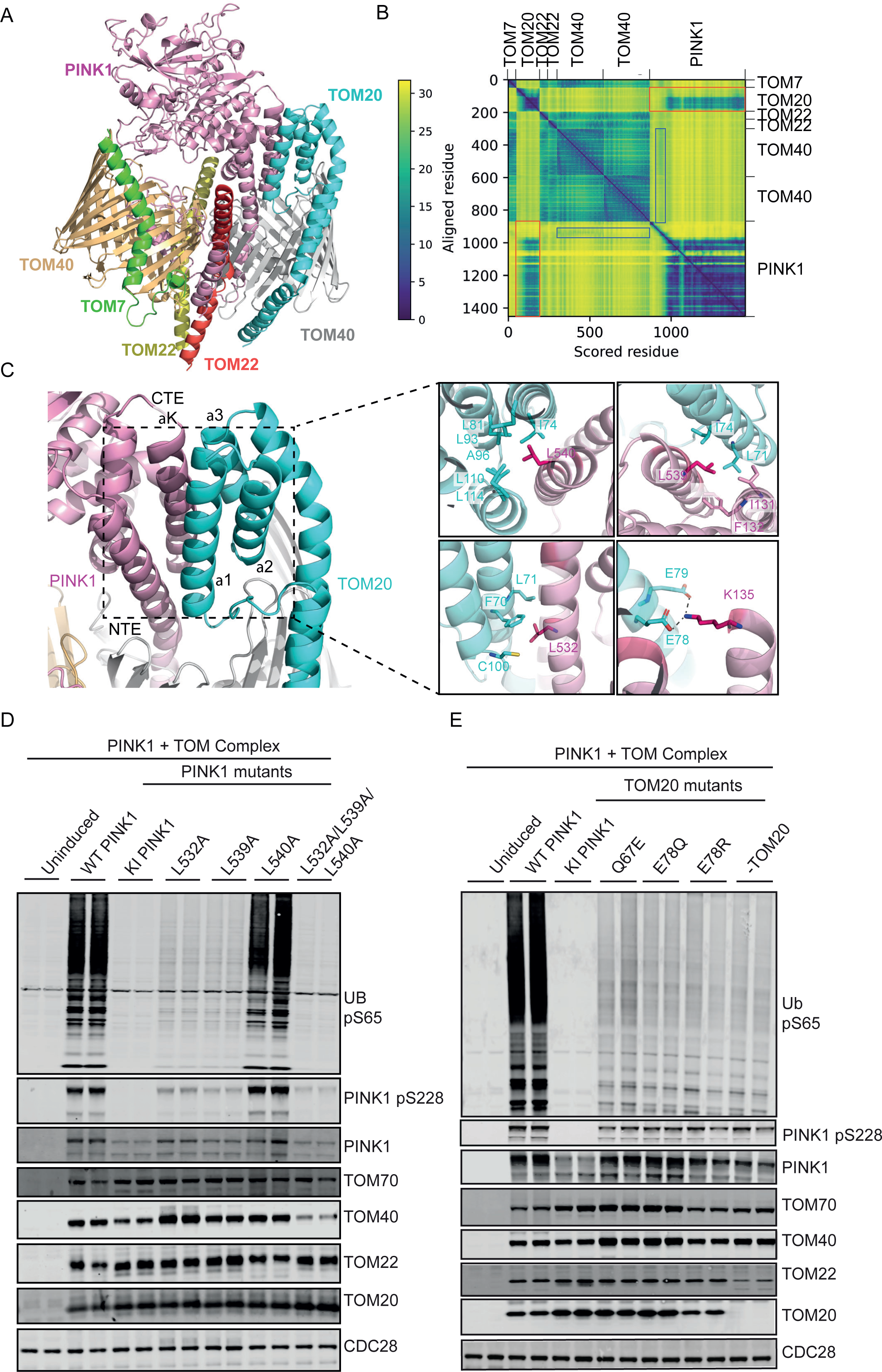
Structural modelling of PINK1-TOM complex predicts direct interaction between PINK1 and TOM20. **(A)** Overall structure of AlphaFold prediction of PINK1 bound to TOM complex (1 molecule TOM7 green; 2 molecules TOM22 yellow/red; dimeric TOM40 orange/grey) indicates direct PINK1 (pink) interaction with TOM20 (cyan). **(B)** Predicted Aligned Error (PAE) plot highlights predicted interaction between PINK1 and TOM20, marked by red boxes. N-terminal segment of PINK1 transverses one TOM40 pore (moderate confidence (blue boxes)), while other TOM components form high confident model as indicated. **(C)** A close-up view illustrates binding interface between PINK1 and TOM20, wherein the N-terminal extension (NTE) and C-terminal extension (CTE) regions of PINK1 interact with C-terminal region of TOM20 (α1-3 helices). Key interactions between conserved amino acids of PINK1 and TOM20 are indicated. **(D-E)** Mutational analysis in yeast cells confirms critical role of PINK1 NTE/CTE interaction with TOM20 for PINK1 activation. **D)**. Hydrophobic leucines on PINK1 at the interface were mutated to alanine (L532A, L539A, L540A and L532A/L539A/L540A), cells expressing these mutants were grown on YP medium supplemented with 2% raffinose, protein expression was induced by the addition of galactose. Cells were harvested lysed, and 20 µg of whole cells lysate was subjected to immunoblot analysis, phospho-ubiquitin was blotted as a readout of PINK1 activation and activity. The membranes were also blotted using the indicated antibodies using Li-COR Odyssey CLx imaging system. **(E)** Residues on TOM20 (Q67 and E78) were also mutated, immunoblot shows significant effect of these mutations on PINK1 activity. Cells with mutant TOM20 were grown on YP medium supplemented with 2% raffinose, protein expression was induced by the addition of galactose. Cells were harvested, processed, and phospho-ubiquitin was blotted as a readout of PINK1 activation and activity. The membranes were also blotted using the indicated antibodies using Li-COR Odyssey CLx imaging system.

All five models predicted direct interaction between the NTE:CTE interface of PINK1 and residues within the C-terminal half of TOM20 (Fig. 3A-C, fig. S6A-B). TOM20 is comprised of an N-terminal transmembrane domain anchoring it to the OMM and the C-terminal region (residues 50-145) that spans five α-helices (α1 - α5) exposed to the cytosol where it binds mitochondrial precursor proteins via their MTS (fig. S7A). Several key conserved residues within the C-terminus of TOM20, namely Gln67, Glu78, Glu79, Phe70, Leu71, Ile74 and Val109, have been found to play a critical role in the recognition and binding of mitochondrial precursor proteins (fig. S7A-B) [45, 46]. Inspection of the AlphaFold PINK1-TOM complex model revealed a major hydrophobic interface between the αK helix of the CTE region of PINK1 and the hydrophobic patch formed by TOM20 α1 and α3 helices comprising multiple highly conserved residues including Leu532, Leu539, and Leu540 of PINK1 CTE and Phe70, Leu71, Ile74, Leu81, Leu110, and Leu114 of TOM20 (Fig. 3C and fig. S7A-E). Furthermore, the NTE of PINK1 formed polar interactions with TOM20 including between the conserved Lys135 residue of PINK1 NTE and Gln78 located at the periphery of the α1 helix of TOM20 (Fig. 3C and fig. S7A-E).

To investigate the functional impact of mutations of these residues, we generated yeast strains in which we expressed PINK1 CTE mutants, L532A, L539A, L540A and a combinatorial L532A/L539A/L540A triple mutant (CTE 3A) together with all TOM complex subunits (Fig. 3D). Immunoblotting analysis of total PINK1 did not reveal any differences in PINK1 processing suggesting that these mutations do not impact on mitochondrial import (Fig. 3D). Single CTE mutants led to mild-to-moderate effects on PINK1 activation but strikingly PINK1 activation was completely abolished in the CTE 3A mutant and this was also associated with a reduction in PINK1 stabilisation (Fig. 3D). In addition, the AlphaFold model predicted electrostatic and hydrogen bond interactions between the NTE region of PINK1 and TOM20 including a predicted salt bridge between the conserved Lys135 of PINK1 and Glu78 of TOM20 and also predicted hydrogen bonding between Gln67 of TOM20 and PINK1 (Fig. 3C and fig. S7A-B). We initially generated a yeast strain in which we expressed the PINK1 NTE mutants, K135E and K135M mutants in complex with all intact TOM complex subunits and immunoblot analysis revealed reduction in ubiquitin phosphorylation consistent with reduced activation (fig. S8). We next generated strains of wild-type PINK1 co-expressed with all TOM complex subunits but in which we expressed TOM20 mutants Q67E, E78Q or E78R (Fig. 3E). Immunoblot analysis revealed that these mutations in TOM20 led to reduced levels of ubiquitin phosphorylation to a similar degree as the minus TOM20 strain (Fig. 3E). Overall, these studies suggest that the PINK1 NTE:CTE interface promotes binding to TOM20.

### Molecular basis of PINK1 interaction with TOM70

We further investigated how PINK1 activation is regulated by TOM70. TOM70 consists of multiple repeating units known as Tetratricopeptide repeat (TPR) domains [47] (fig. S9A). The N-terminal TPRs form a loosely structured region called the NTD-pocket, which primarily interacts with heat shock proteins [48, 49] (fig. S9A). The C-terminal TPRs form the CTD-pocket which specifically binds to mitochondrial preproteins for import into the mitochondria [50] (fig. S9A). We employed the ColabFold notebook [42] to run AlphaFold [43] structure predictions of the PINK1 with TOM70, imputing full-length human PINK1 and TOM70 sequences. AMBER structure relaxation was used to ensure appropriate orientation of the side chains to avoid steric clashes. Five models of a PINK1-TOM70 complex were generated from higher to lower confidence based on inter-PAE values (Fig. 4A-B and fig. S10A-B). Three of the five models predicted a highly consistent binding interface between an N-terminal region of PINK1 (residues 71-106) and a region located within the CTD of TOM70 (Fig. 4C and fig. S10A-B) with high PAE scores for interacting regions (fig. S10A-B). Previous studies have suggested that PINK1 contains an internal MTS sequence in this region that acts redundantly with the canonical N-terminal MTS [39, 51]. The Hermann lab previously elaborated an algorithm to predict internal MTS-like signals for TOM70 in the mature region [52]. We imputed PINK1 into this algorithm (https://csb-imlp.bio.rptu.de) and this predicted several MTS-like regions of which the strongest peak mapped to the TOM70-binding region predicted by AlphaFold (fig. S10C). This predicted interface represents an extensive binding surface with multiple contacts involving hydrogen bonding, hydrophobic interactions, and electrostatic interactions. Electrostatic surface mapping visualisation of TOM70 demonstrated that the CTD pocket was largely negatively charged and the corresponding N-terminal interacting surface region of PINK1 was positively charged (fig. S10D). Furthermore, a series of conserved positively charged residues in PINK1 (Arg 83, Arg88, and Arg 98) are predicted to form salt bridges with a series of conserved negatively charged residues within TOM70 (Asp488, Asp545, Glu549 and Asp229 respectively) (Fig 4C). To test the predicted mode of binding model, we generated yeast strains expressing PINK1 mutations in the TOM70 interface (R83A/R88A/R98A triple mutant; R83E/R88E/R98E triple mutant). We observed that the combined triple mutation largely abolished ubiquitin phosphorylation (Fig. 4D). We next generated strains expressing TOM70 mutations within the PINK1-TOM70 binding interface namely D488A, D545A, and E549A and all of these substantively lowered ubiquitin phosphorylation to a similar degree as the minus TOM70 strain (Fig. 4E; fig. S9B-D). Overall, our data indicate that TOM70 is also required for optimal stabilisation and activation of PINK1 at the TOM complex via an internal MTS-like region that we define as the TOM70 interacting region (TIR).

**Figure 4.**
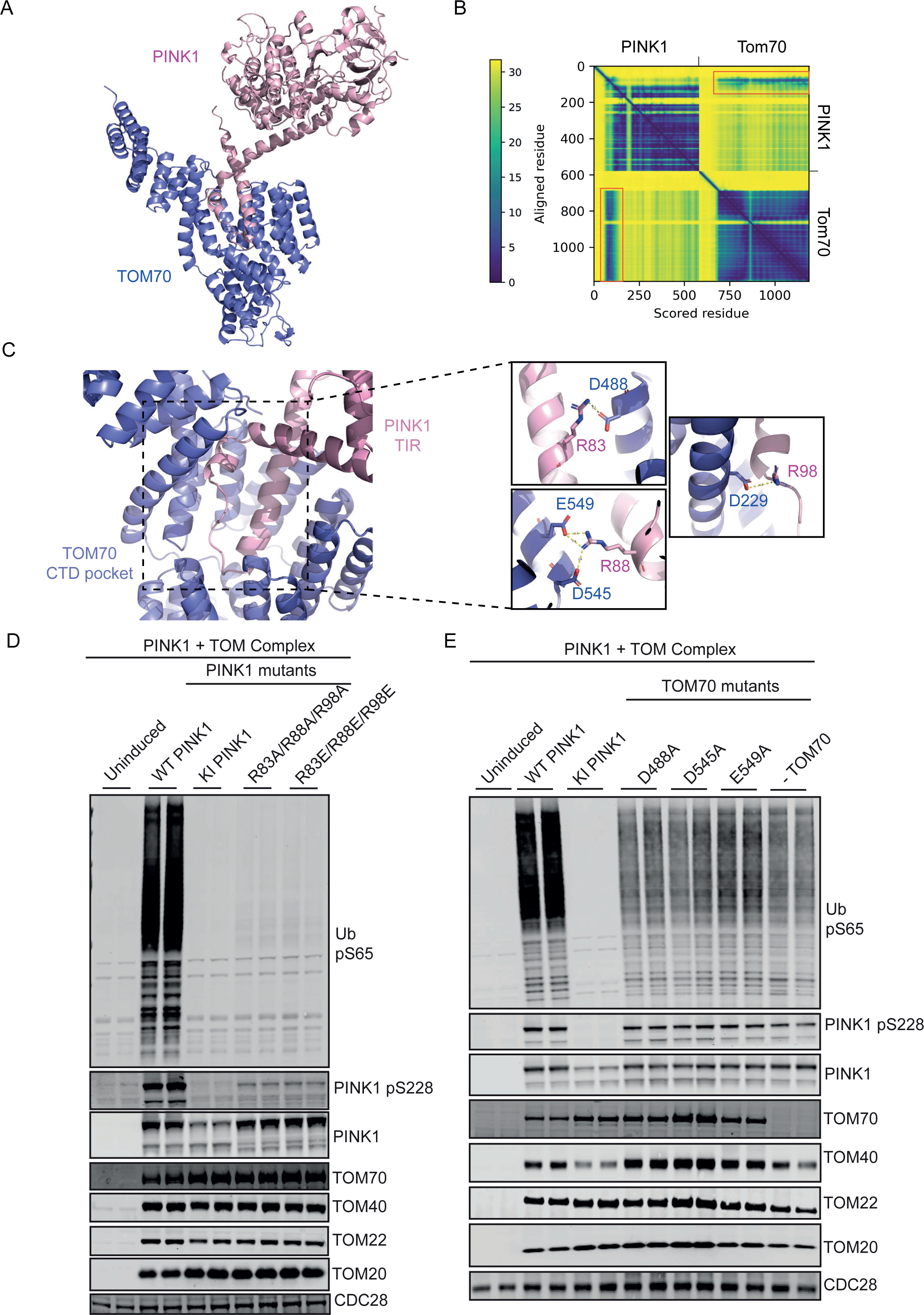
PINK1 binds TOM70 with N-terminal region preceding NTE (residues, 60-111). **(A)** AlphaFold model of PINK1 in complex with TOM70. PINK1 is coloured in pink and TOM70 in purple. **(B)** Predicted Aligned Error (PAE) plot highlights the interaction between PINK1 and TOM70, indicated with red boxes. **(C)** Close-up view shows residues making direct interaction between the two proteins. **(D)** The three arginine residues (R83, R88 and R98) on PINK1 making interactions with TOM70 were mutated and the effect on PINK1 activity were compared with wild type PINK1 and Kinase inactive PINK1. Residues on TOM70 (D488, D545 and E549) making interactions with PINK1 were mutated to alanine and the effect of these mutations on PINK1 activity was assayed and compared with the wild type, kinase inactive and the minus TOM70 cells. Cells carrying these mutations and the corresponding controls were grown on YP medium supplemented with 2% raffinose, protein expression was induced by the addition of galactose. Cells were harvested, processed and 21hosphor-ubiquitin was blotted as a readout of PINK1 activation and activity. The membranes were also blotted using the indicated antibodies using Li-COR Odyssey CLx imaging system.

### Validation of predicted PINK1-TOM interactions in mammalian cells

To validate the PINK1-TOM20 predicted interaction in a mammalian cell system, we generated stable cell lines in which we re-introduced full-length wild-type PINK1-3FLAG (WT); kinase-inactive mutant PINK1 (KI); CTE PINK1 mutants namely L532A, L539A, L540A and a combined CTE 3A mutant into Flp-In T-REx HeLa PINK1-knockout cells (generated by exon 2-targeted CRISPR-Cas9 [13]. To determine the effect of the selected PINK1 mutants on activation, cells were treated with DMSO or 10 μM Antimycin A / 1μM Oligomycin (A/O) for 3 h to induce mitochondrial depolarization (Fig. 5A). Immunoblot analysis of whole cell extracts revealed that CTE single point mutants led to a slight reduction of phosphorylated ubiquitin upon mitochondrial depolarization of which L540A had a more noticeable effect, however, consistent with the analysis in yeast, phosphorylated ubiquitin, was completely abolished in the CTE 3A mutant (Fig. 5B-C). We next investigated the PINK1-TOM70 interaction and generated stable cell lines in which we re-introduced PINK1 TIR mutants namely R83A, R88A, R98A and combined R83A/R88A/R98A or R83E/R88E/R98E triple mutants into Flp-In T-REx HeLa PINK1-knockout cells (Fig. 5D-E). Immunoblot analysis of whole cell extracts revealed that TIR single point mutants led to minimal reduction of phosphorylated ubiquitin upon mitochondrial depolarization, however, phosphorylated ubiquitin, was most reduced in the TIR 3A or 3E mutant (Fig. 5E). These data indicate that the PINK1:TOM interfaces identified in our yeast system are also functionally relevant in mammalian cells.

**Figure 5.**
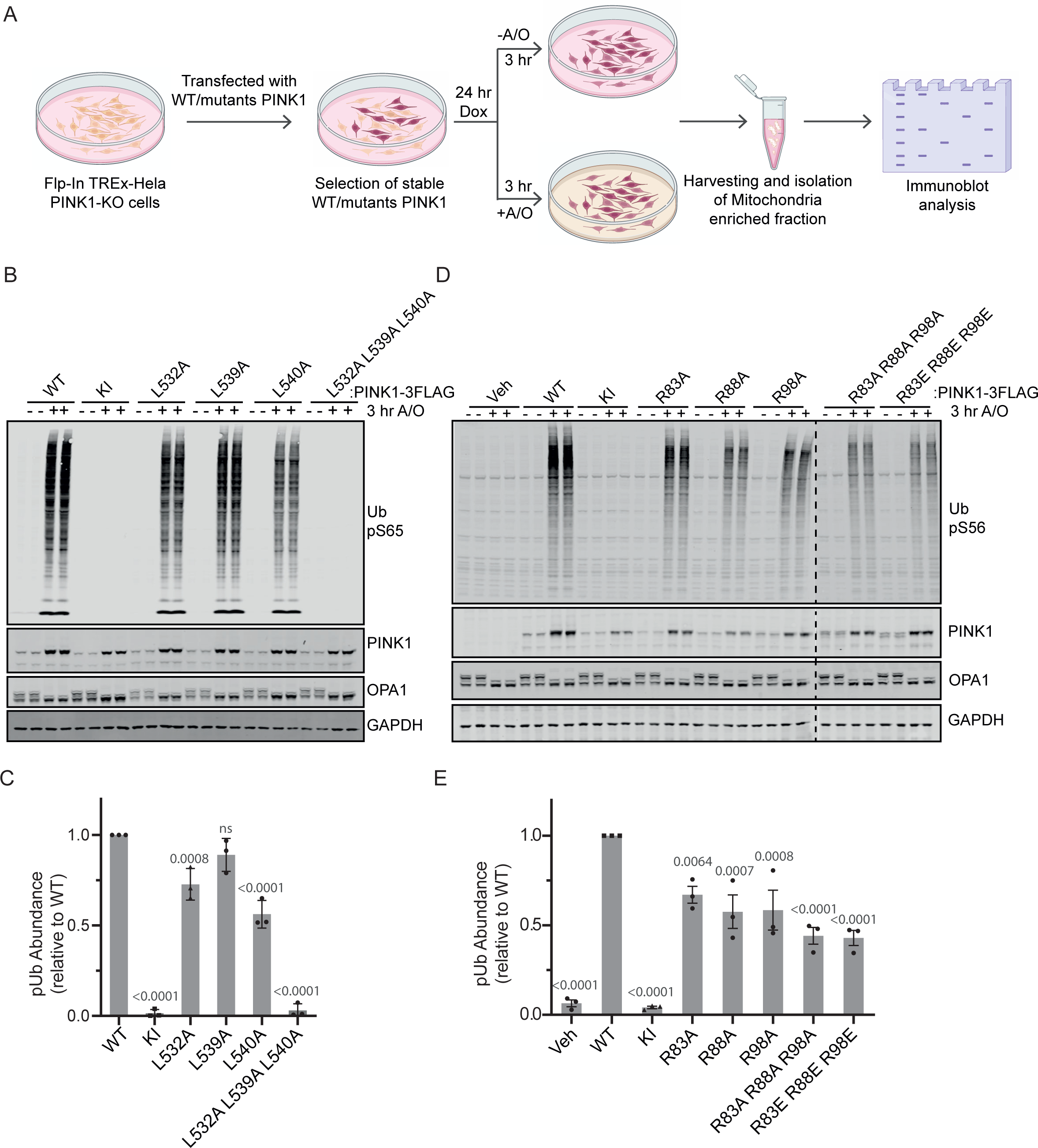
Optimal PINK1 activation requires interaction with TOM20 and TOM70 in mammalian cells following mitochondrial depolarisation. **(A)** Schematic of workflow of analysis of PINK1 TOM-binding mutants in PINK1 knockout Flp-In Trex HeLa cells. The schematic image was made using BioRender. **(B)** PINK1 CTE TOM20-defective binding mutants lead to reduced PINK1 activation. Stably expressing PINK1-3FLAG WT, KI (D384A), and CTE mutant (L532A, L539A, L540A and triple (L532A/L539A/L540A)) cell lines were generated in PINK1-knockout Flp-In TRex HeLa cells. PINK1-3FLAG expression was induced by 24 h treatment with 0.2 μM doxycycline, and mitochondrial depolarization induced by 3 h treatment with 10 μM Antimycin A / 1 μM Oligomycin (A/O) where indicated. Whole cell lysates were subjected to immunoblotting with anti-PINK1 (in-house/DCP antibody), anti-Ub pS65 (CST), anti-OPA1 (BD) and anti-GAPDH primary antibodies. Data representative of 3 independent experiments. **(C)** Quantification for immunoblots of analysis of CTE mutants (L532A, L539A, L540A and triple (L532A/L539A/L540A)) were quantified for Ub pS65 (pUb) relative to WT PINK1 as mean ± SD (n = 3). Statistical analysis was done by ordinary one way ANOVA where p values relative to WT are shown above the bars. **(D)** PINK1 TOM70-defective binding mutants lead to reduced PINK1 activation. Stably expressing PINK1-3FLAG WT, KI (D384A), and N-terminal mutant (R83A, R88A, L98A and triple (R83A/R88A/L98A or R83E/R88E/R98E)) cell lines were generated in PINK1 knockout Flp-In TRex HeLa cells. PINK1-3FLAG expression was induced by 24 h treatment with 0.2 μM doxycycline, and mitochondrial depolarization induced by 3 h treatment with 10 μM Antimycin A / 1 μM Oligomycin (A/O) where indicated. Whole cell lysates were subjected to immunoblotting with anti-PINK1 (in-house/DCP antibody), anti-Ub pS65 (CST), anti-OPA1 (BD) and anti-GAPDH primary antibodies. Data representative of 3 independent experiments. **(E)** Quantification of immunoblots for analysis of CTE mutants (R83A, R88A, L98A and triple (R83A/R88A/L98A or R83E/R88E/R98E)) were quantified for Ub pS65 (pUb) relative to WT PINK1 as mean ± s.d. (n = 3). Statistical analysis was done by ordinary one-way ANOVA where p values relative to WT are shown above the bars.

We further investigated the role of TOM20 and TOM70 in stabilising PINK1 at the TOM complex of mammalian cells. We performed BN-PAGE analysis in HeLa cells stably expressing wild-type PINK1-3FLAG and observed two complexes consistent with previous studies [13, 14, 26, 27]. Immunoblotting analysis of these complexes indicated that TOM20, and TOM40 were present in the native ∼500 kDa TOM complex and the ∼700kDa PINK1-TOM complex whereas TOM70 was not detectable in either of the complexes (fig. S11A-D). PINK1 resided in two HMW complexes namely the ∼700kDa PINK1-TOM complex and a complex of intermediate size between the ∼500 kDa and ∼700kDa complexes as previously reported [13, 14, 26, 27] (fig. S11A). We next performed BN-PAGE analysis on stable cell lines in which we re-introduced full-length wild-type PINK1-3FLAG (WT); kinase-inactive mutant PINK1 (KI); a combined CTE L532A/L539A/L540A mutant (TOM20-binding deficient); and a combined TIR R83E/R88E/R98E mutant (TOM70-binding deficient) into Flp-In T-REx HeLa PINK1-knockout cells (Fig. 6A). Strikingly immunoblotting analysis with anti-TOM40 antibodies demonstrated that both the CTE L532A/L539A/L540A and TIR R83E/R88E/R98E triple mutants prevented the formation of the ∼700kDa PINK1-TOM complex and this was also confirmed by immunoblotting with anti-PINK1 antibody (Fig. 6B). Overall, these studies demonstrate that PINK1 recruitment, stabilisation, and activation at the TOM complex requires interaction with TOM20 and TOM70 in mammalian cells and supports our findings from the yeast reconstitution system.

**Figure 6.**
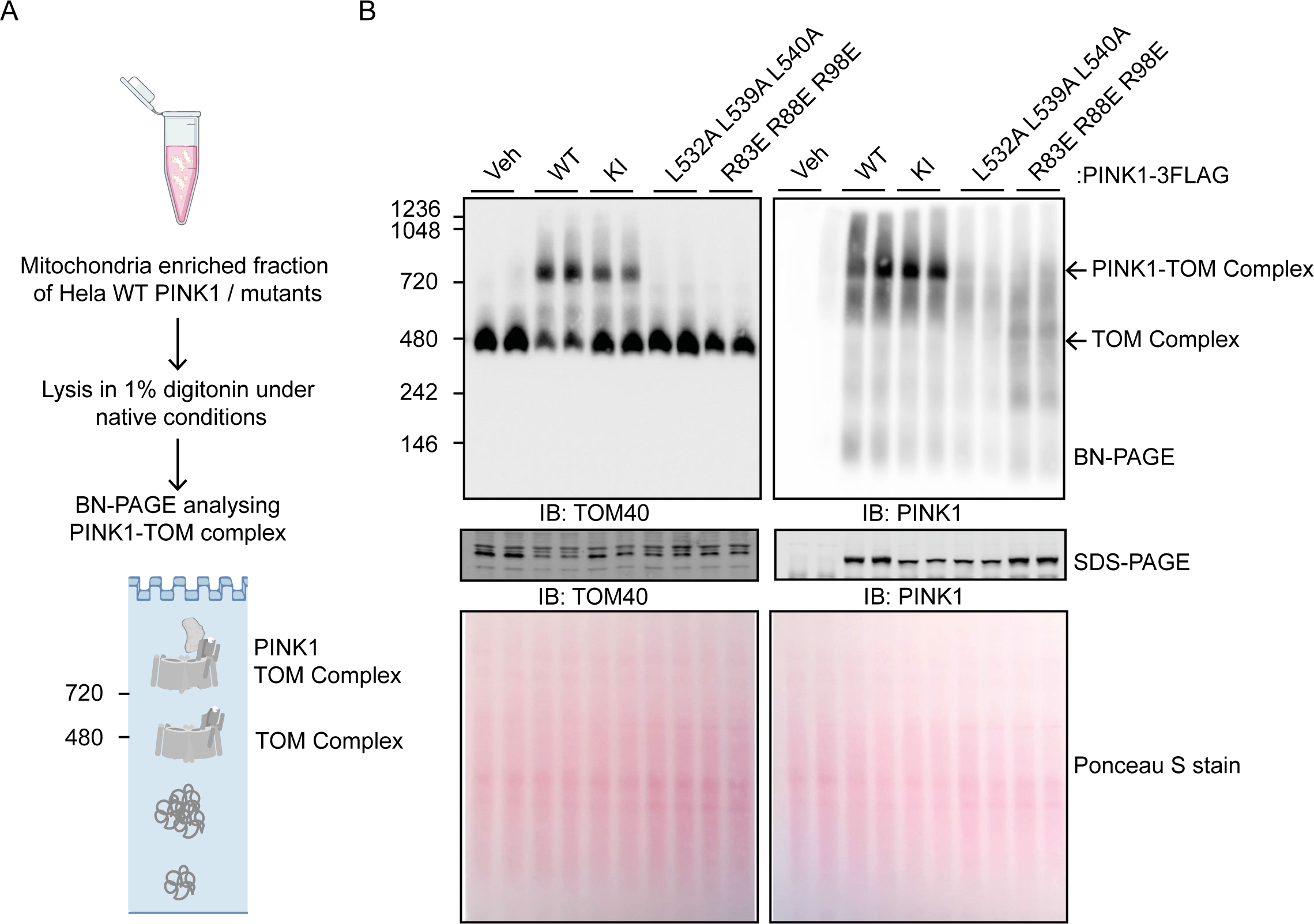
PINK1 stabilisation at the TOM complex is dependent on TOM20 and TOM70 interaction in mammalian cells. **(A)** Schematic of workflow of BN-PAGE analysis of PINK1 TOM-binding mutants in HeLa cells. **(B)** Mitochondrial enriched fractions were isolated from PINK1 knockout Flp-In TRex HeLa cells stably expressing PINK1-3FLAG WT, KI (D384A), TOM20-defective binding mutant (L532A/L539A/L540A) and TOM70-defective binding mutant (R83E/R88E/R98E) mutants were treated with A/O for 3 h and then subjected to BN-PAGE and immunoblotted for anti-TOM40 or anti-PINK1 antibodies. Samples were also subjected to SDS-PAGE and immunoblotting with anti-TOM40 or anti-PINK1 antibodies, and total protein visualized by Ponceau S staining.

## DISCUSSION

Our yeast reconstitution system enables genetic dissection of the contribution of human TOM complex subunits towards human PINK1 activation. These findings highlight the essential requirement of the TOM40 pore-containing core complex as well as a role for the TOM20 and TOM70 receptor subunits for optimal PINK1 activation (Fig. 2 & Fig. 7).

**Figure 7.**
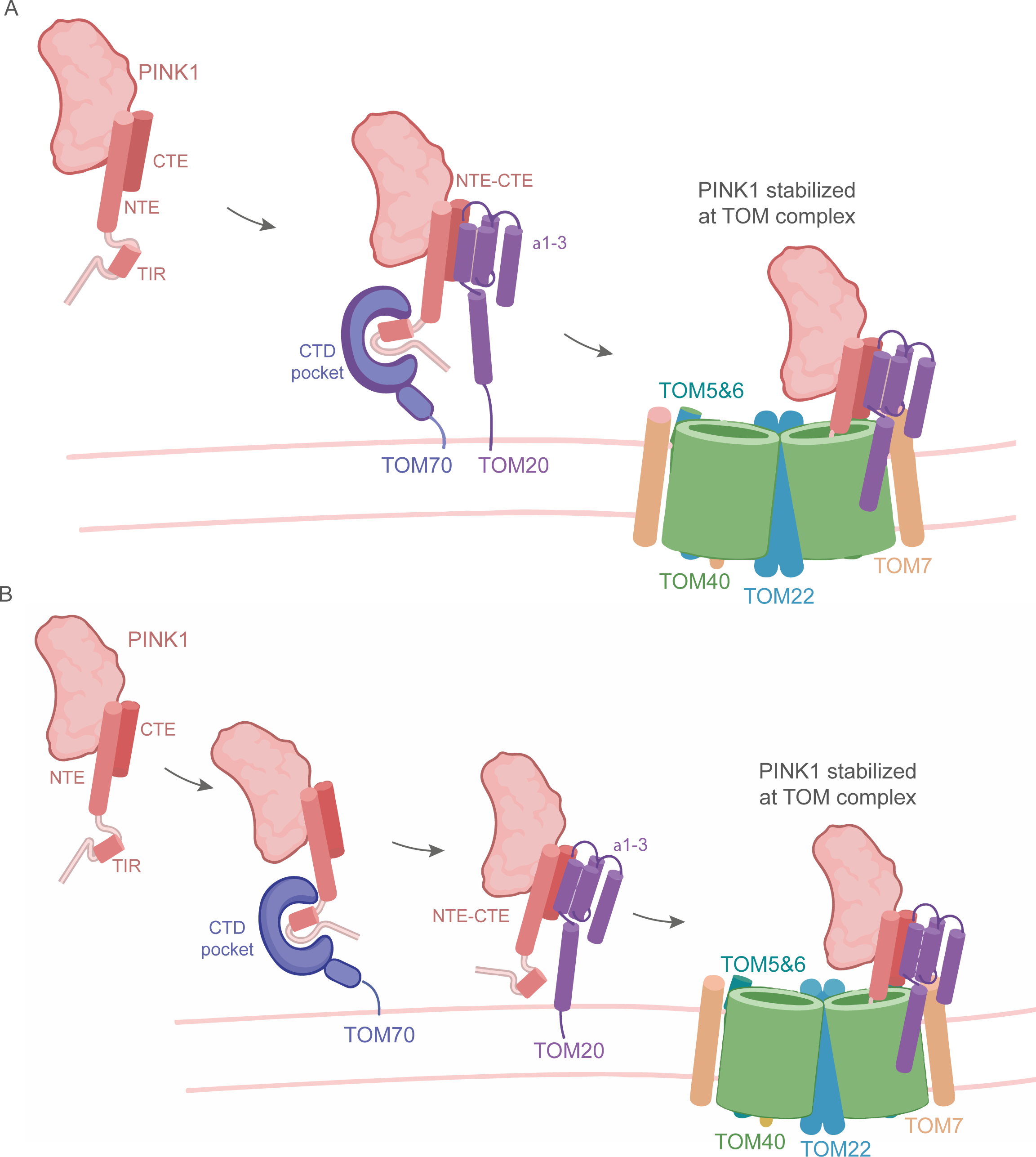
Schematic models of role of TOM70 and TOM20 interaction with PINK1 mediating stabilisation and activation at the TOM complex at sites of damaged mitochondria. **(A)**. Interaction of the PINK1 TIR region with the CTD pocket of TOM70 occurs concurrently with interaction of the PINK1 NTE:CTE interface with the C-terminal α1-3 helices of TOM20 and is required for PINK1 stabilisation at the TOM complex. Image was created using BioRender.com **(B)** Sequential binding of PINK1 TIR region to CTD pocket of TOM70 followed by PINK1 NTE:CTE interface with the C-terminal α1-3 helices of TOM20 is required for PINK1 stabilisation at the TOM complex. Abbreviations: TOM70 interacting region (TIR); N-terminal extension (NTE); C-termimal extension (CTE); C-terminal domain (CTD); Translocase of outer membrane (TOM). Image was created using BioRender.com

TOM20 typically binds mitochondrial proteins via N-terminal mitochondrial targeting sequences (MTS) consisting of 15-40 amino acids [45, 46, 53–57] that form an amphipathic helix [58]. Previous analysis of the PINK1 MTS sequence (aa 1-34) showed that it is sufficient for mitochondrial targeting to the matrix in polarised mitochondria of mammalian cells [59–61]. Following mitochondrial depolarisation, the stabilisation of PINK1 at the TOM complex is distinct compared to other MTS-containing mitochondrial proteins which are generally released into the cytosol and not stabilised at the TOM complex. A role for TOM20 in PINK1 stabilisation upon mitochondrial depolarisation was first reported using cross-linking/immunoprecipitation studies of PINK1 under denaturing conditions and showing that only TOM20 could be co-isolated with PINK1 under these conditions suggesting a direct interaction [26]. However, the mechanism remained elusive since full-length PINK1 lacking the MTS could still be imported to mitochondria and stabilised and activated on depolarised mitochondria [39, 51]. We and others recently defined the NTE:CTE interface as being critical for PINK1 recruitment to the TOM complex [12–14] and our new findings support a model whereby the NTE:CTE interface of active PINK1 binds to the C-terminal cytosolic domain of TOM20 (Figs. 2-3; Fig.7). The cytosolic domain of TOM20 has previously been shown to have a diverse selectivity for client mitochondrial proteins mediated by a hydrophobic MTS sequence motif that is extremely broad (*φXXφφ; φ* is hydrophobic acid and *x* is any other amino acid) [45]. Structural studies have mapped the TOM20 peptide binding site to a hydrophobic pocket within its cytosolic domain (e.g.[45]). Similar to MTS presequences, the binding of PINK1 CTE to TOM20 is majorly directed by hydrophobic leucine residues although the sequence of the PINK1 hydrophobic patch (aa 532 – LQQSAATLL – aa 540) displays notable differences from the general MTS sequence motif (fig. S7E). A further notable difference is that polar interactions between TOM20 and PINK1 are directed by residues located within the distinct NTE α-helix whereas these typically occur at the periphery of MTS presequences [45] (Fig 3C). Overall the interaction between PINK1 and TOM20 that we have identified further expands the diversity of TOM20 client interactions (Fig. 3 & Fig. 7).

In future work it will be interesting to understand how TOM20 switches from binding the PINK1 MTS under polarised mitochondria conditions to the NTE:CTE interface in depolarised mitochondria. This is likely to be due to the flexibility of the cytosolic domain of TOM20 which renders it highly dynamic and accounts for it not being visible on previous high resolution cryo-EM structures of the TOM complex [33–35]. Recently, low-resolution cryo-EM structures of the TOM complex in *Neurospora crassa* have revealed multiple sub-complexes containing TOM20, TOM22 and TOM40 in differing stoichiometries including one TOM20 in a dimeric TOM40 complex [36]. This suggests that TOM20 is able to adopt myriad conformations and interactions within the overall complex [36]. AlphaFold modelling predicted an asymmetric PINK1-TOM complex with 1 molecule of PINK1 bound to 1 molecule of TOM20 within a dimeric TOM40 complex (Fig. 3), however, in view of the flexibility of TOM20 we cannot rule out additional subcomplexes of PINK1 and TOM20 with the rest of the TOM components with varying stoichiometries (Fig. 3). Furthermore, the *Neurospora crassa* structure revealed a strong interaction of the acidic patch of the N terminus of TOM22 with a positively charged patch in cytosolic TOM20 helix suggesting that TOM20 requires to dock with TOM22 for optimal receptor function [36], which was also not observed in the AlphaFold model.

A previous study of PINK1 had mapped an outer mitochondrial localisation signal (OMS) (residues 70 – 94) at its N-terminus located between the MTS and transmembrane domain (TMD) [51]. This region is dispensable for mitochondrial import but is required for PINK1 stabilisation and activation at the TOM complex on depolarised mitochondria in a TOM40 dependent manner [39, 51]. TOM70 has been reported to recognise internal signals in hydrophobic regions and consistent with this was shown to be required for PINK1 import [62] however, a role for TOM70 in PINK1 activation in depolarised mitochondria was lacking since BN-PAGE studies reported that TOM70 was not detectable in the ∼700 kDa PINK1-TOM complex [26, 27]. It was recently reported that TOM70 acts as the major receptor for PINK1 import and stabilisation in depolarised mitochondria based on siRNA knockdown studies of TOM subunits in mammalian cells [49]. Our genetic dissection of TOM subunits in the reconstitution assay provides a clearcut role of TOM70 for PINK1 activation (Fig. 2 & Fig. 4) and we have mapped a TOM70 interaction region (TIR) that contains overlapping residues with the previously elaborated OMS region. AlphaFold modelling predicted binding of the N-terminal region of PINK1 to the C-terminal cytosolic domain of TOM70 (Fig. 4). Our results contrast with a previous study that employed peptide mapping studies to define the TOM70 binding site on PINK1 and suggested that TOM70 can interact with the N-terminal MTS, OMS or TMD regions with equivalent affinity [49]. To date there is no experimental structure of mammalian TOM70 but crystal structures of yeast TOM70 revealed it contains 26 α-helices that are involved in the formation of 11 tetratricopeptide repeat (TPR) motifs [63]. The N-terminal domain functions to bind chaperones including Hsp90 in mammals [48, 63, 64]. Binding sites for selected mitochondrial preproteins have been mapped to the C-terminus (α-helices 8-26) although to date a general receptor site/motif has yet to be defined [50]. The crystal structure of yeast TOM70 has revealed a highly conserved groove located in the centre of the C-terminal domain which has been attributed to be the major binding site for mitochondrial preproteins (TPR 4-11) [63]. The distal side of the binding groove is made of hydrophobic and polar residues whereas the proximal side contains three highly conserved negatively charged residues Glu473, Glu542 and Glu577 [63]. AlphaFold modelling suggests PINK1 forms specific interactions with the proximal residues and also has interactions with the distal side that may aid in interaction by acting as docking site as it has been suggested for other preproteins [50]. Previous studies have mapped numerous Parkin-mediated ubiquitylation sites in the TOM70 C-terminus (K473, K504, K539, K573 and K607) [65, 66], however, these are located on the outside of the C-terminus and not likely to affect the binding surface (fig. S9). Similarly regulatory phosphorylation sites on the C-terminus of TOM70 have been reported [67] it will be interesting to assess the role of these modifications as well as chaperone binding to the N-terminus on the interaction between PINK1 and TOM70 under depolarised mitochondrial conditions.

In contrast to a previous study [49], our studies unambiguously show that both TOM70 and TOM20 are important for PINK1 stabilisation and activation at the TOM complex (Fig. 7). AlphaFold was unable to predict a high confidence model with both TOM70 and TOM20 in the PINK1-TOM complex suggesting that PINK1 interactions with these subunits is likely to be dynamic and this is compounded by dynamic interactions of the cytosolic domains of TOM20 and TOM70 with other subunits of the TOM complex if bound to PINK1 simultaneously. We propose a model that PINK1 is engaged by both TOM20 and TOM70 for recruitment and stabilisation at the TOM complex (Fig, 7A), however, we cannot exclude that the association of PINK1 with TOM70 and TOM20 subunits occurs in a sequential manner and since TOM70 is not detectable in the PINK1-TOM complex on BN-PAGE (fig. S11D), suggesting that it may bind to PINK1 earlier to direct it to the TOM20 subunit and core complex (Fig. 7B).

Our reconstitution analysis confirms the essential role of the TOM7 subunit for PINK1 activation as first revealed by the Youle lab [38, 39]. AlphaFold modelling did not predict a direct interaction between TOM7 and PINK1 and instead was in line with previous Cryo-EM analysis showing that the three small TOM subunits, 5, 6 and 7 are peripherally bound to TOM40 via distinct interactions [33–35]. We did not observe much effect of loss of TOM5 or TOM6 on activation whereas loss of TOM7 largely abolished PINK1 activation akin to loss of TOM22 that is critical for stabilising the TOM40 dimeric pore structure (Fig. 2). Whilst the role of the small TOMs in the mammalian TOM complex are still to be fully elucidated [30], our findings are in keeping with previous analysis that have shown a critical role for TOM7 (but not TOM5 and 6) in maintaining stability of the mammalian TOM40 core complex [37].

An unexpected and striking finding from our studies was that co-expression of human PINK1 with TOM subunits was sufficient for its activation (Fig. 1B-C). Over the past decade, PINK1 activation has mainly been studied in the context of mitophagic signalling following mitochondrial depolarisation, induced by mitochondrial uncouplers (e.g. AO or CCCP), or mitochondrial matrix misfolding stress, triggered by matrix Hsp90 inhibitors or over-expression of the deletion mutant of OTC [3–5]. Since all lead to PINK1 activation, our reconstitution results suggest that in the context of mitochondrial damage, the key event for PINK1 activation is its stabilisation at the TOM complex. Previous studies have shown weak activation of PINK1 when it is transiently over-expressed in mammalian cells without any mitochondrial damage [41], and we speculate the low activity is due to the substoichiometric levels of the TOMs compared to over-expressed PINK1. Thus, it would be interesting to assess whether this activity is increased if PINK1 is co-transfected with exogenous human TOM subunits. Similarly we detected weak activation of full-length human PINK1 when expressed alone in yeast or in yeast strains in which human TOM20 or TOM70 were mutated to prevent PINK1 interaction (Fig. 3E & 4E) and this may be due to interaction with the yeast TOM20 and/or TOM70 since the PINK1 interaction sites are conserved (fig. 7B & fig. 9B). Previous studies have reported that recombinant expression of catalytic domain-containing fragments of human PINK1 in *E. coli* or insect cells display no significant catalytic activity associated with low expression yields and unstable protein [40]. In contrast, recombinantly expressed catalytic domain-containing fragments of insect orthologues of PINK1 (e.g. Pediculus humanus corporis) are active with high expression yields of monodisperse protein [40]. In our hands we have been unable to identify any N-terminal or C-terminal truncated His-tagged human PINK1 constructs, out of >30 tested, that when expressed in insect cells lead to high yields of soluble human PINK1 as detected by colloidal Coomassie gels (fig. S12A-D). Furthermore, replacement of the Ins1 loop of human PINK1 with the orthologous region of Pediculus PINK1 or the pseudokinase PEAK1 [68] (fig. S12A-B), did not lead to enhanced expression of soluble protein (fig. S12D). Collectively our findings would suggest that recombinant co-expression of TOM subunits with human PINK1 in insect cells is needed to generate high yields of stable PINK1 and reconstitute PINK1 activation *in vitro*.

In future work, experimental structural data is required to better understand PINK1 regulation within the TOM complex which, although challenging due to transient association of TOM20 and TOM70 as well as PINK1, may provide additional molecular insights into PINK1 activation including dimerization which was recently shown for insect PINK1 and how this mediates Ser228 transphosphorylation [12, 14]. In addition to Ser228, we recently identified Ser167 autophosphorylation as being important for activation of human PINK1 which is not conserved in insect PINK1 [69]. Furthermore, several critical residues on PINK1 NTE/CTE and TIR that bind TOM20 (fig. S7D-E) and TOM70 (fig. S9D) respectively are also not conserved in the insect orthologues suggesting there will be important differences in active human PINK1 from previously solved insect PINK1 structures. Overall, our current analysis provides new insights into human PINK1 activation that will be of utility in the development of small molecule activators as a therapeutic strategy against PD.

## MATERIALS AND METHODS

### Molecular Biology and Cloning

For mammalian expression, PINK1-3FLAG constructs were cloned into pcDNA5 vectors for recombination into HEK293 Flp-In TRex Hela PINK1 knockout cell lines [13]. For the yeast strains, codon optimised plasmids to express human PINK1-3FLAG, TOM70, TOM40, TOM22, TOM20, TOM7, TOM6 and TOM5 were generated, using previously described methods [70]. Human genes were paired into four genetically modified yeast vectors (pORs), based on the pRS vector series that enables efficient shuttling of vectors in specific yeast strains for facile manipulation [71] (fig. S1). yOR1 strain harbours human PINK1-3FLAG + TOM40; yOR2 strain harbours PINK1-3FLAG + TOM40 + TOM22 + TOM7; yOR3 strain harbours PINK1-3FLAG + TOM40 + TOM22 + TOM7 + TOM70 + TOM20; and yOR4 strain harbours PINK1-3FLAG + TOM40 + TOM22 + TOM7 + TOM70 + TOM20 + TOM5 + TOM6 (Supplementary Table S1 (Table. S1)). The paired genes were cloned either side of the GAL1_10 promoter which allows the bidirectional induction of both genes upon addition of galactose to the growth medium. Site-directed mutagenesis was carried out using the QuikChange method with KOD polymerase (Novagen). All yeast and mammalian constructs (Supplementary Table S2 (Table. S2)) were verified by The Sequencing Services (School of Life Sciences, University of Dundee) and are now available to request via the MRC PPU Reagents and Services website (https://mrcppureagents.dundee.ac.uk/). For cloning of hPINK1 constructs for insect cell expression, the coding sequences for the PINK1 constructs were PCR amplified using clone OHu25380D (Genscript) as a template and cloned into the vector pFB-6HZB (SGC) as previously described [72]. Expression from this vector yields proteins with a TEV protease-cleavable N-terminal His6-Z tag.

### Antibodies for biochemical analysis

The following antibodies were used in this study: α-FLAG (Sigma), α-HSP60 (CST), α-OPA1 (BD biosciences, 612606), α-PINK1 (Novus, BC100-494), α-PINK1 (DCP), α-ACSL1 (Cell signalling technologies, 4047S), α-ubiquitin pSer65 (Cell signalling technologies, 37642), α-TOM20 (Santa Cruz); α-TOM22 (Abcam), α-TOM40 (Abcam); α-TOM70 (Aviva Systems Biology), CDC28(Santa cruz-Sc6709). We sought to generate in-house antibodies against TOMs 5, 6 and 7 due to lack of available commercial antibodies. Successful generation of TOM6 (fig. S13A-B) and TOM7 (fig. S13C-D) antibodies was confirmed by testing using whole cell lysates from HEK 293 cells, yeast whole cell lysates and mitochondria fraction from yeast cells. TOM5 antibody generation was not successful (data not shown). TOM6 and TOM7 antibodies are available from Reagent and services, School of Life Sciences, University of Dundee. The polyclonal α-PINK1 pSer228 antibody was generated by the Michael J. Fox Foundation’s research tools program in partnership with Abcam (Development of a monoclonal antibody is underway. Please contact for questions.) All fluorophore-conjugated mouse, rabbit, and sheep secondary antibodies for immunoblotting and immunofluorescence were obtained from Sigma.

### Expression of human PINK1 and TOM complex in yeast cells

Codon optimised plasmids to express the eight subunits of human PINK1-TOM complex from the bidirectional *GAL1*, *10* promoters in budding yeast were generated, using previously described methods [70]. The Saccharomyces cerevisiae strain YCE1164 (MATa ade2-1 ura3-1 his3-11,15 trp1-1 leu2-3,112 can1-100 bar1Δ::hphNT pep4Δ::ADE2) was transformed with linearised plasmids using standard procedures to generate protein expression strains. For the protein expression strains, the codon usage of the synthetic gene constructs was optimised for high-level expression in *Saccharomyces cerevisiae*, as described previously [70].

### Yeast protein induction and expression

300 µl of overnight culture from positive clones were inoculated into 10 ml of fresh YP medium supplemented with 2 % raffinose and then grown at 30°C with 180rpm shaking, to an OD600 of 1.7. Protein expression was subsequently induced by adding galactose to a final concentration of 2 % and continuing growth for a further 10h. Cells were harvested by centrifugation at 3000xg for 10min, flash frozen in liquid nitrogen and stored until needed.

### Yeast protein extraction and quantification

To extract proteins, frozen cells were allowed to thaw on ice and resuspended in 200 µl 20% Trichloroacetic acid (TCA), lysed by vortexing for 35 sec in the presence of glass beads. After the beads settle, 100 µl of the supernatant was collected into a fresh microfuge tube. An additional 200 µl of 5% TCA was added to the beads and vortexed for a further 35 sec, and 150 µl of the supernatant was also collected after allowing the beads settle and added to the previous supernatant collection. The supernatant was centrifugated at maximum speed at 4°C in a microfuge to collect the precipitated protein, which was then resuspended in 200 µl of 25 mM Tris, 300 mM NaCl, 10% glycerol and 0.5 mM TCEP pH 7.5 containing protease inhibitor cocktail. Protein quantification was done using BCA method and bovine serum albumin as standard (BSA).

### Immunoblotting

20 μg of protein was subjected to SDS–PAGE (4–12% Bis-Tris gel) and transferred onto nitrocellulose membranes. Membranes were blocked for 1 h in Tris-buffered saline with 0.1% Tween (TBST) containing 5% (w/v) milk. Membranes were then probed with the indicated antibodies in TBST containing 5% (w/v) BSA overnight on a roller at 4°C. Detection was performed using appropriate secondary antibodies and scanned using Li-COR Odyssey CLx imaging system.

### Isolation of yeast mitochondria

Yeast mitochondria were purified following the protocol outlined by Gregg et al. (2009). In brief, a 2L culture of yeast cells carrying wildtype PINK1 alone or wildtype PINK1/KI with all TOMs was cultured, and expression was induced as described. Cells were harvested by centrifugation at 3000xg for 10 minutes at room temperature. The resulting pellet was washed twice with 1.2 M Sorbitol and resuspended in DTT buffer (100 mM Tris-H_2_SO_4_ pH 9.4, 10 mM DTT) at a ratio of 2 ml of buffer per gram of cells. The suspension was gently rotated at 70rpm at 30°C for 30 minutes. After a subsequent centrifugation and resuspension in Zymolase buffer (20 mM K3PO4 pH 7.4, 1.2M sorbitol), the cells were treated with Zymolase powder (1 mg/gm of wet cells) and rotated gently at 70rpm for 60 minutes at 30°C. The resulting spheroplasts were centrifuged for 5 minutes at 3000g at 4°C. Maintaining a cool temperature, the spheroplast was then resuspended in ice-cold homogenization buffer (10 mM Tris-HCl pH 7.4, 0.6M sorbitol, 1mM EDTA, 0.2% BSA) and transferred into a pre-cooled ice glass homogenizer. Using a tight pestle, homogenization was performed with 15 strokes. Following homogenization, differential centrifugation steps at 2000×g and 3000×g were performed to discard cell debris. The isolated mitochondria were then centrifuged at 15,000×g, and to enhance purity, the mitochondria were resuspended in SEM buffer (10 mM MOPs/KOH pH7.2 250 mM sucrose, 1 mM EDTA buffer) and subjected to another centrifugation step. The final mitochondria were resuspended in SEM buffer, and their protein concentration was adjusted to 10 mg/ml using a Bradford protein assay.

### AlphaFold modelling

To gain insight into how PINK1 might interact with the TOM complex, or how PINK1 might interact with TOM70, AlphaFold prediction tool was deployed [73]. AMBER structure relaxation was used to ensure appropriate orientation of the side chains to avoid steric clashes. The resulting output models were ranked by confidence level and analysed by visualization using PyMol. All key interactions are listed in Supplementary Table 3 (Table. S3).

### Blue native PAGE (BN-PAGE)

The samples for blue native PAGE (BN-PAGE) analysis were prepared using a Native PAGE Sample Prep Kit (Invitrogen). For BN-PAGE, mitochondria-enriched fractions were gently pipetted up and down 10 times in 1x Native PAGE buffer with 1% digitonin followed by an incubation for 30 min at 4°C. The samples were centrifuged at 20,000*g* for 30 min at 4°C. Samples were quantified by BCA assay and supplemented with 0.002% G-250 (Invitrogen). BN-PAGE was performed by Native PAGE Running Buffers (Invitrogen). The gels were transferred on to PVDF membranes for IB analysis. PVDF membranes were washed in 100% methanol and subjected to immunoblotting.

### Mammalian cell culture and transfection

HeLa wild-type (WT) and PINK1 knockout cells were routinely cultured in standard DMEM (Dulbecco’s modified Eagle’s medium) supplemented with 10% FBS (fetal bovine serum), 2 mM L-Glutamine, 100 U ml−1 Penicillin, 100 mg ml−1 Streptomycin (1X Pen/Strep) and 1 X non-essential amino acids (Life Technologies). HeLa Flp-In TREx cells were cultured using DMEM (Dulbecco’s modified Eagle’s medium) supplemented with 10% FBS (foetal bovine serum), 2 mM L-glutamine, 1x Pen/Strep, and 15 μg/ml blasticidin. Culture media was further supplemented with 100 μg/ml zeocin pre-recombination with PINK1-3FLAG constructs. Transfections were performed using the polyethylenimine method. To ensure uniform expression of recombinant proteins, stable cell lines were generated in a doxycycline-inducible manner. HeLa Flp-In TREx CRISPR-mediated PINK1 knockout cells were generated. The PINK1 null host cells containing integrated FRT recombination site sequences and Tet repressor were co-transfected with 4.5/9 µg of pOG44 plasmid (which constitutively expresses the Flp recombinase) and 0.5/1 µg of pcDNA5-FRT/TO vector containing a hygromycin resistance gene for selection of the gene of interest with FLAG tag under the control of a doxycycline-regulated promoter. Cells were selected for hygromycin and blasticidin resistance 3 days after transfection by adding fresh medium supplemented with 15 µg ml−1 of blasticidin and 100 µg ml−1 of hygromycin. Protein expression was induced by the addition of 0.1 μg/ml doxycycline for 24 hours. Mitochondrial depolarisation was induced by treatment with 10 μM AntimycinA and 1 μM Oligomycin (Sigma; prepared in DMSO) for 3 hrs. Cells were harvested and resuspended in mitochondrial fractionation buffer (20mM HEPES pH 7.5, 250 mM sucrose, 3 mM EDTA, 5 mM sodium β-glycerophosphate, 50 mM sodium fluoride, 5 mM sodium pyrophosphate, 1 mM sodium orthovanadate, 200 mM chloracetamide). Cell suspensions were physically disrupted by 25 passes through a 25-gauge needle, and debris removed by centrifugation at 800 x g. The resulting supernatant was subject to centrifugation at 16,600 x g to harvest a mitochondria-enriched pellet. Samples for SDS-PAGE were generated from mitochondrial lysates resulting from resuspension in mitochondrial fractionation buffer with 1% Triton.

### Insect cell expression

Cloning of hPINK1 constructs for insect cell expression and test purifications The coding sequences for the PINK1 constructs were PCR amplified using clone OHu25380D (Genscript) as a template and cloned into the vector pFB-6HZB (SGC) as previously described [72]. Expression from this vector yields in proteins with a TEV protease-cleavable N-terminal His6-Z tag. Baculoviruses were then generated according to protocols from the Bac-to-Bac expression system (Invitrogen). For protein expression, exponentially growing TriEx cells (3 mL suspension, 2×10^6^ cells/mL, Novagen) in serum-free Insect-XPRESS medium (Lonza) were infected with recombinant virus (MOI>2) and cultured for 66 hours at 27°C under gentle agitation. Cells were harvested by centrifugation (20 min, 1000x g, 4°C), resuspended in lysis buffer (50 mM HEPES pH 7.4, 500 mM NaCl, 0.5 mM TCEP, 5% glycerol, either with or without 0.05% digitonin) and lysed via sonication (24-tip horn, 35% amplitude, 5 s pulse / 10 s pause, 3 min total pulse time). The lysate was cleared by another round of centrifugation (30 min, 13,000 rpm, 4°C) and loaded onto 25 µL pre-equilibrated Ni-NTA beads (#17526802, Cytiva) in gravity flow columns. After washing the beads with lysis buffer, His6-Z-PINK1 was eluted in lysis buffer containing 300 mM imidazole. Samples of the total lysate and elution were analyzed side-by-side in CriterionTM T Precast gels (Bio-Rad). Gels were stained with Coomassie or further processed for Western blotting and immunodetection using anti-PINK1 (#BC100-494, Novus Biologicals) or anti-hexahistidine antibody (#SAB2702220, Sigma-Aldrich). A detailed description of all tested constructs is given in Supplementary Table S4 (Table. S4).

### Live cell imaging

Yeast colonies grown on respective plates were inoculated into YP liquid media with 2% raffinose as carbon source and incubated overnight at 30°C. On the following day, cells were diluted, grown to exponential phase (OD600=0.3-0.6) and protein expression was initiated by the addition of 2% galactose for a total of 3 h 45min before completion of the incubation period, mitochondria were stained by addition of 500 nM of MitoTracker CMXRos Red. At the end of the incubation time, cells were briefly spun down, washed 2x with PBS and applied to Concavalin A-coated coverslips which were then placed on glass-slides for immediate image acquisition using a Leica DMi8. Further processing was carried out in the Leica LAS X software platform which includes histogram adjustment and denoising with THUNDER (Leica).

### Statistical Analysis

Statistical analysis was done by ordinary one-way ANOVA using GraphPad Prism 9.5.1. Dunnett’s multiple comparisons test was performed with a single pooled variance relative to WT. p values relative to WT are shown as stars above the bars; p≤ 0.05, 0.01, 0.001, 0.0001 are represented as *, **, ***, **** respectively.

## ACKNOWLEDGEMENTS

We thank Yogesh Kulathu for helpful advice on AlphaFold structural modelling. We also thank Kyunglin Min and Eunyong Park for discussions on PINK1 and TOM complex analysis. We express our thanks to Thomas Macartney and the late Mark Peggie for molecular biology and cloning. We thank Nicole Polinski and Shalini Padmanabhan at the Michael J Fox Foundation for advice on PINK1 tool development. We are grateful to the sequencing service (School of Life Sciences, University of Dundee); James Hastie for expression and generation of recombinant proteins (MRC PPU); the MRC PPU tissue culture team (co-ordinated by Edwin Allen) and MRC PPU Reagents and Services antibody teams (co-ordinated by James Hastie). This work was supported by a Wellcome Trust Senior Research Fellowship in Clinical Science (210753/Z/18/Z to M.M.K.M.); the Michael J. Fox Foundation (M.M.K.M.), EMBO YIP Award (M.M.K.M.), the Medical Research Council (core grant MC UU 12016/13 to K.L.), Cancer Research UK (Programme Grant C578/A24558 and PhD studentship C578/A25669 to K.L.). S.M., N.R., R.F.-B. and M.M.K.M. acknowledge funding from The Michael J. Fox Foundation for Parkinson’s Research (MJFF) through grant MJFF-010458. S.M., R.F.-B. and M.M.K.M. were funded by the joint efforts of The MJFF and the Aligning Science Across Parkinson’s (ASAP) initiative. MJFF administers grants ASAP-000519, ASAP-000463, ASAP-000282 and on behalf of ASAP and itself. R.F.-B. acknowledges funding from the Deutsche Forschungsgemeinschaft (DFG, German Research Foundation) through Germany’s Excellence Strategy—EXC2067/1—390729940.

## Conflict of Interest

M.M.K.M. is a member of the Scientific Advisory Board of Montara Therapeutics Inc.

---

**Supplementary Figure 1. Experimental design and constructs. (A)** Schematic depiction of sequential yeast transformation with plasmids carrying two components of the complex each. After each transformation, positive clones were selected on an appropriate nutrient dropout agar plate. The positive clone was then used for the next transformation, this continues until cells carrying all eight components of the complex (PINK1 and all the TOMs) was made. Schematic diagram was made using BioRender.com. **(B)** Schematic depiction of the constructs boundaries for all the proteins used in the experiment. All proteins are full length with flag tag at the C-terminus of PINK1 for detection.

**Supplementary Figure 2. Clonal effect on PINK1 activation. (A)** Immunoblot of four independent clones. Four independent clones were selected separately for wild-type PINK1 with all TOMs, kinase inactive PINK1 with all TOMs and cells expressing PINK1 alone. Cells were grown in similar conditions and treated the same. After protein expression cells were harvested, lysed, and analysed by immunoblot. Interestingly similar results were obtained for all clones. Wildtype PINK1 with all TOMs shows significant level of phospho-ubiquitin compared with uninduced cells and with cells expressing PINK1 alone. As expected, cells expressing kinase inactive PINK1 did not show any activity as compared with the wild type. **(B)** Quantification of phospho-ubiquitin abundance between cells expressing wildtype PINK1+ all TOMs, kinase inactive PINK1+all TOMs and wildtype PINK1 alone. Data represent four independent clones. **(C)** Quantification of PINK1 abundance between cells expressing wildtype PINK1+ all TOMs, kinase inactive PINK1+all TOMs and wildtype PINK1 alone. Data represent four independent clones. **(D).** Quantification of PINK pS228 abundance between cells expressing wildtype PINK1+ all TOMs, kinase inactive PINK1+all TOMs and wildtype PINK1 alone. Data represent four independent clones.

**Supplementary Figure 3. Effect of Parkinson’s disease mutations on kinase activity of PINK1 expressed in yeast in company of the TOM complex. (A)** Wild-type cells, kinase inactive cells (KI) and the mutant cells (Q126P, E240K, G309D and 534_535InsQ) were grown on YP medium supplemented with 2% raffinose, protein expression was induced by the addition galactose to 2% final concentration. Cells were harvested, processed, and phospho-ubiquitin was blotted as a readout of PINK1 activation and activity. The membranes were also blotted using the indicated antibodies using Li-COR Odyssey CLx imaging system. All mutants show significant reduction in phospho-ubiquitin level. Although PINK1 level was slightly reduced compared to wildtype, the level of pS228 was significantly reduced compared with the wildtype except for in G309D cells which is slightly increased compared to the other mutants. **(B)** PINK1 and the human TOMs expressed in the yeast are localised to the yeast mitochondria. Mitochondria was isolated and solubilised using 1% digitonin from cells expressing wildtype PINK1 + the TOMs (WT+TOM complex) and kinase inactive PINK1+ the TOMs (KI+TOM Complex). 10 µg of samples was run on a gel and transferred onto a nitrocellulose membrane; the membrane was then blotted using the indicated antibodies.

**Supplementary Figure 4. Investigation of clonal effect of TOM subunit (TOMs 5, 6, 7, 20 and 22) elimination on PINK1 activation and activity.** Four colonies each were selected after transformation of cells expressing PINK1 and the TOMs minus each indicated subunits. Along with cells containing wild type PINK1 as well as Kinase inactive PINK1 and all the TOMs subunits, cells were grown on YP medium supplemented with 2% raffinose, protein expression was induced by the addition galactose to 2% final concentration. Cells were harvested, processed, and phospho-ubiquitin was blotted as a readout of PINK1 activation and activity. Interestingly, all clones tested showed similar pattern with slight variation in minus TOM5 clones. The membranes were also blotted using the indicated antibodies using Li-COR Odyssey CLx imaging system.

**Supplementary Figure 5. Investigation of clonal effect of TOM subunit (TOMs 40, 70 and 20/70) elimination on PINK1 activation and activity.** Four colonies each were selected after transformation of cells expressing PINK1 and TOMs minus each indicated subunits. Along with cells containing the wild type PINK1 as well as Kinase inactive PINK1 and all the TOMs subunits, cells were grown on YP medium supplemented with 2% raffinose, protein expression was induced by the addition galactose to 2% final concentration. Cells were harvested, processed, and phospho-ubiquitin was blotted as a readout of PINK1 activation and activity. Interestingly all clones tested showed similar pattern. The membranes were also blotted using the indicated antibodies using Li-COR Odyssey CLx imaging system.

**Supplementary Figure 6. Robust PINK1-TOM complex Interaction from AlphaFold-Generated Models. (A)** Detailed PINK1-TOM complex is shown with pLDDT mapping on the structure on the right. **(B)** Five distinct models of PINK1-TOM complex were generated using locally installed AlphaFold. All five models consistently depict a robust interaction between PINK1 and TOM20 with high confidence. Despite improper folding of the TOM40 pores in Model 5, the PINK1-TOM20 interaction remained consistent. **(C)** The PAE plots of respective models depicted in A above showing strong interaction between PINK1 and TOM20.

**Supplementary Figure 7. Multiple Sequence Alignment of PINK1 and TOM20. (A)** Schematic representation of TOM20 highlighting its five alpha helices. PINK1-binding regions are highlighted with shaded red boxes. **(B)** Multiple sequence alignment of TOM20, generated using MUSCLE and visualized in Jalview. Residues highlighted by black arrows signify those associated with the loss of PINK1 activity in the yeast reconstitution system. Red arrows indicate conserved residues known to participate in hydrophobic interactions and are also involved in the interaction with the hydrophobic residues of PINK1. **(C)** Schematic of PINK1 and domain outline. **(D)** Multiple sequence alignment of the NTE region of PINK1, with a schematic of PINK1 above the sequence. Residues mutated are highlighted by arrows **(E)** Multiple sequence alignment of the CTE region of PINK1, generated using MUSCLE and visualized in Jalview. Residues mutated are highlighted with arrows.

**Supplementary Figure 8. K135 mutation at the PINK1-TOM20 interface have significant effect on PINK1 activity.** Lysine135 making hydrogen bond interactions with Glu78 and Glu79 was mutated to K135E and K135M respectively, cells expressing these mutants were grown on YP medium supplemented with 2% raffinose, protein expression was induced by the addition of galactose. Cells were harvested lysed, and 20 µg of whole cells lysate was subjected to immunoblot analysis, phospho-ubiquitin was blotted as a readout of PINK1 activation and activity. The membranes were also blotted using the indicated antibodies using Li-COR Odyssey CLx imaging system. The K135E activity reduced 2.3-fold compared to the wildtype while the K135M mutation decreased PINK1 activity further with about 7.5-fold.

**Supplementary Figure 9. Multiple Sequence Alignment of PINK1 and TOM70. (A)** Schematic representation of TOM70 highlighting TPR binding region in the N-terminal domain and C-terminal domain. PINK1-binding region are highlighted with shaded red boxes. **(B)** Multiple sequence alignment of TOM70, generated using MUSCLE and visualized in Jalview. Residues highlighted by black arrows signify those associated with the loss of PINK1 activity in the yeast reconstitution system. **(C)** Schematic of PINK1 and domain outline. **(D)** Multiple sequence alignment of the of the TOM70 interaction region (TIR) of PINK1, generated using MUSCLE and visualized in Jalview. Residues mutated are highlighted by black arrows.

**Supplementary Figure 10. PINK1-TOM70 AlphaFold-Generated Models**. **(A)** Five distinct models of PINK1-TOM70 interactions were generated using locally installed AlphaFold. Three models consistently depict a robust interaction between PINK1 and TOM70 with high confidence. Model 4 predicts interaction between MTS and TOM70 whilst Model 5 showed no interaction. **(B)** The PAE plots of respective models depicted in A above showing strong interaction between PINK1 and TOM70. **(C)** A propensity map for Internal Mitochondrial Targeting-like sequences (iMTS-L) was generated using the TargetP prediction tool. The prediction revealed the highest peak in a region analogous to that predicted by AlphaFold. Additionally, several other regions with lower scores were also identified. **(D)** Electrostatic surface potential mapping, performed using PyMOL, highlighted the negative CTD core of TOM70 and the corresponding positive PINK1 N-terminal region.

**Supplementary Figure 11. PINK1-TOM complex within the mammalian system.** The PINK1-TOM complex in both Vehicle and PINK1-3FLAG stable cell lines were examined using Blue Native Polyacrylamide Gel Electrophoresis (BN-PAGE) with specific antibodies, including **(A)** PINK1 antibody, **(B)** TOM40 antibody, **(C)** TOM20 antibody, and **(D)** TOM70 antibody. The presence of the PINK1-TOM complex was observed using TOM40 and PINK1 antibodies, confirming its association with the TOM complex. Both TOM receptors (TOM20 and 70) were scrutinized within the PINK-TOM complex, however, TOM70 was undetectable in the final PINK1-TOM complex.

**Supplementary Figure 12. Test expression of hPINK1 constructs using the Baculovirus expression system. (A)** The full-length hPINK1 protein was not stable when expressed in insect cells. Several N-terminal and C-terminal truncations were probed for their potential to increase stability. Further, constructs without a probably unfolded insertion (Ins1) were also generated. A detailed description of all tested constructs is given in Table. S4. (**B)** Workflow from cloning, expression to purification testing. Notably, lysis buffers with and without detergents were tested to maximize the amount of soluble hPINK1. (**C)** The total lysates (T) and the eluates from Ni-NTA (E) for every construct were analyzed by SDS-PAGE. Stably expressed constructs are expected to show a distinct band in the E samples. For hPINK1, however, no stably expressed constructs were identified. WB analysis using an anti-PINK1 antibody confirmed that all constructs were expressed, but not enriched during the Ni-NTA purification. (**D)** Purification testing of the Ins1 deletion constructs. None of the proteins tested were enriched during the affinity chromatography.

**Supplementary Figure 13. Generation of TOM6 and TOM7 antibodies**. In-house-generated antibodies TOM6 and TOM7 were analysed for their specificity. For detection in mammalian, the HEK293 cells were transfected with HA-tagged TOM6/7 at either N-terminus or C-terminus. The TOM6/7 were immunoprecipitated by HA-affinity beads and analysed by HA-antibody or TOM6/7 antibody. **(A)** Immunoblot of mammalian cell lysate for TOM6 **(B)** Immunoblot of yeast whole cell lysate and mitochondria fraction for TOM6 **(C)** Immunoblot of mammalian cell lysate for TOM7. **(D)** Immunoblot of yeast whole cell lysate and mitochondria fraction for TOM7.

**Supplementary video**: A movie showing high confidence PINK1-TOM complex and zoom revealing the interface between PINK1 and TOM20. Coloured in pink is PINK1, in orange and grey are TOM40 core, in cyan is TOM20, in yellow and red is TOM22 and in green is TOM7.

## Notes

### Competing Interest Statement

Miratul Muqit is a member of the Scientific Advisory Board of Montara Therapeutics Inc.

### Summary of Updates

correction of nomenclature for yeast strains used addition of a further Supplementary Table correction of Figure 3

