## Supplementary Figures for "Mechanism of human PINK1 activation at the TOM complex in a reconstituted system"

Supplementary Figure 1

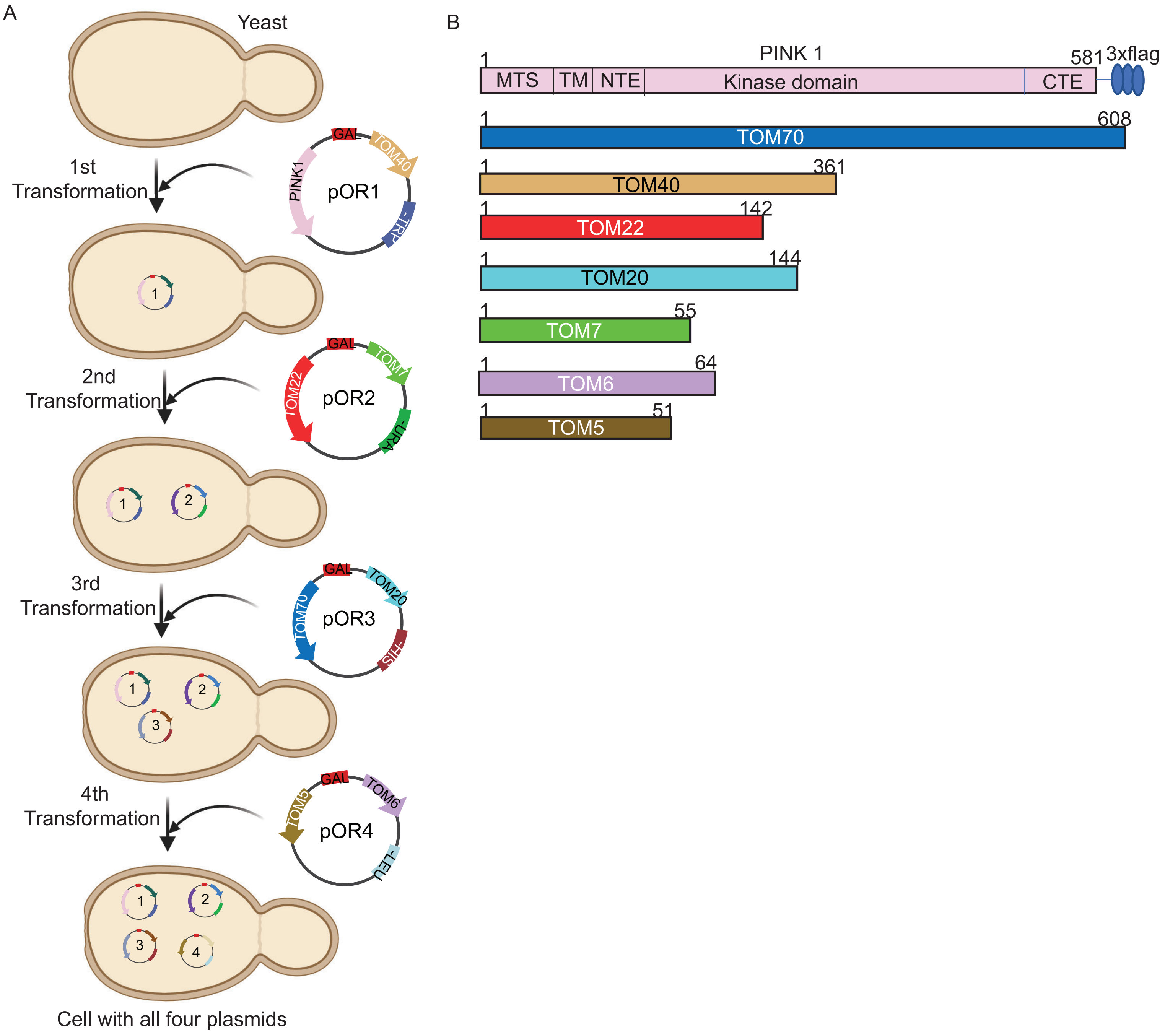

Supplementary Figure 2

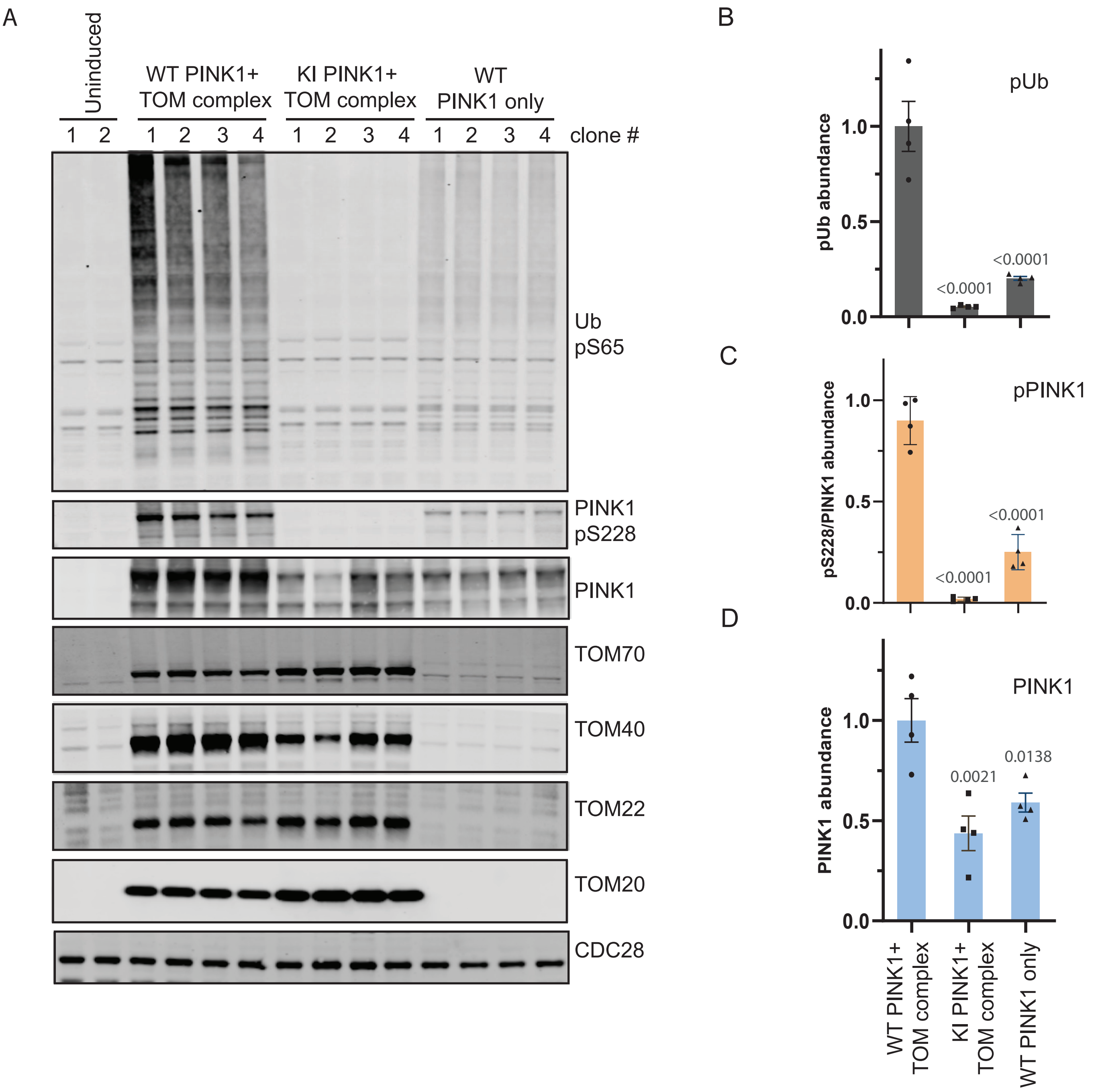

Supplementary Figure 3

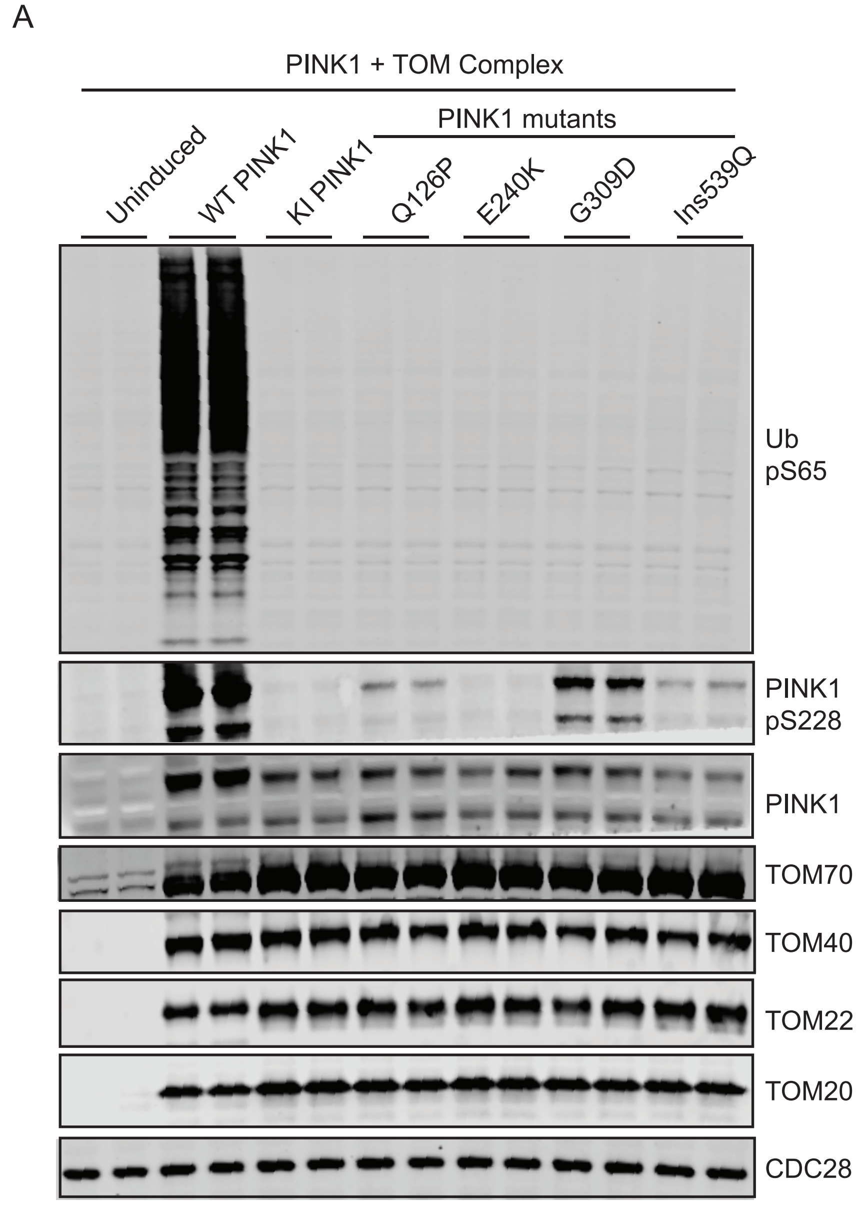


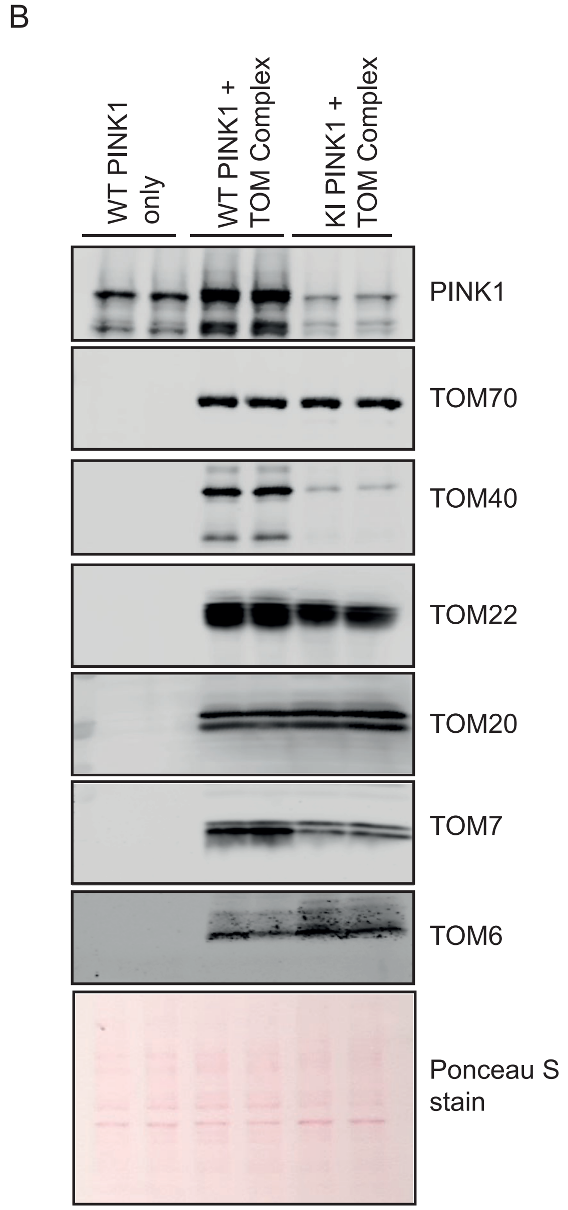

Supplementary Figure 4

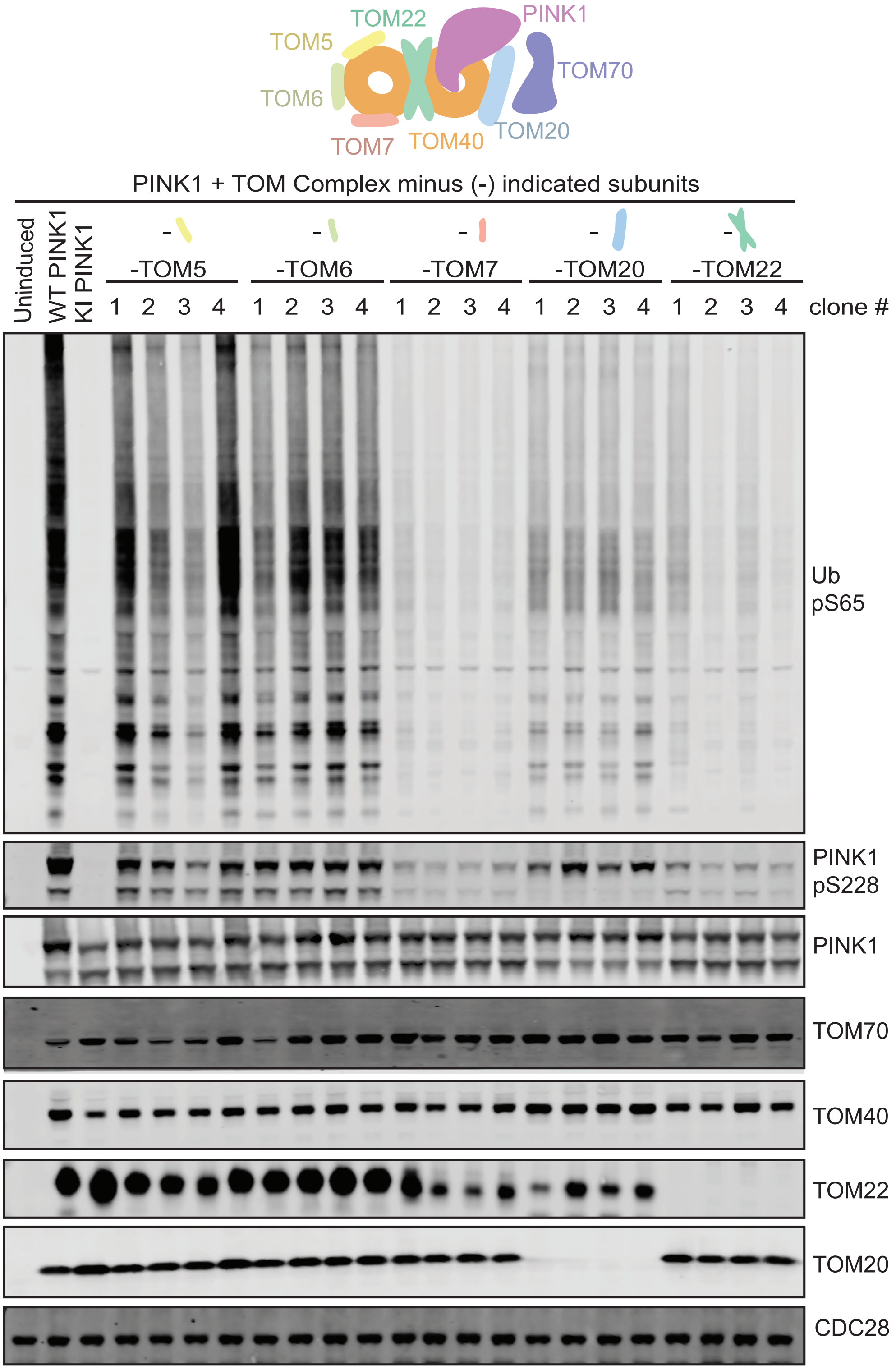

Supplementary Figure 5

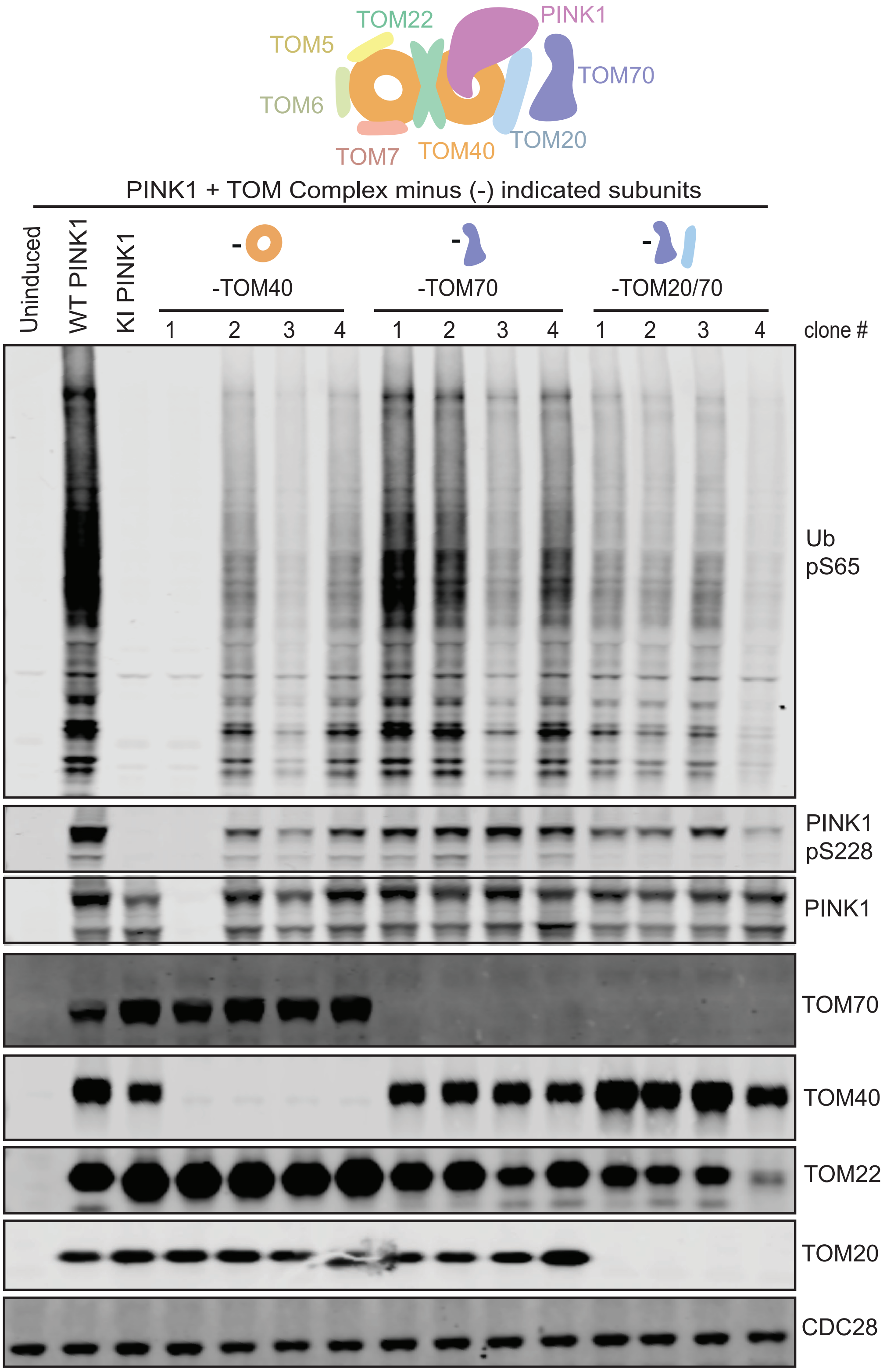

Supplementary Figure 6

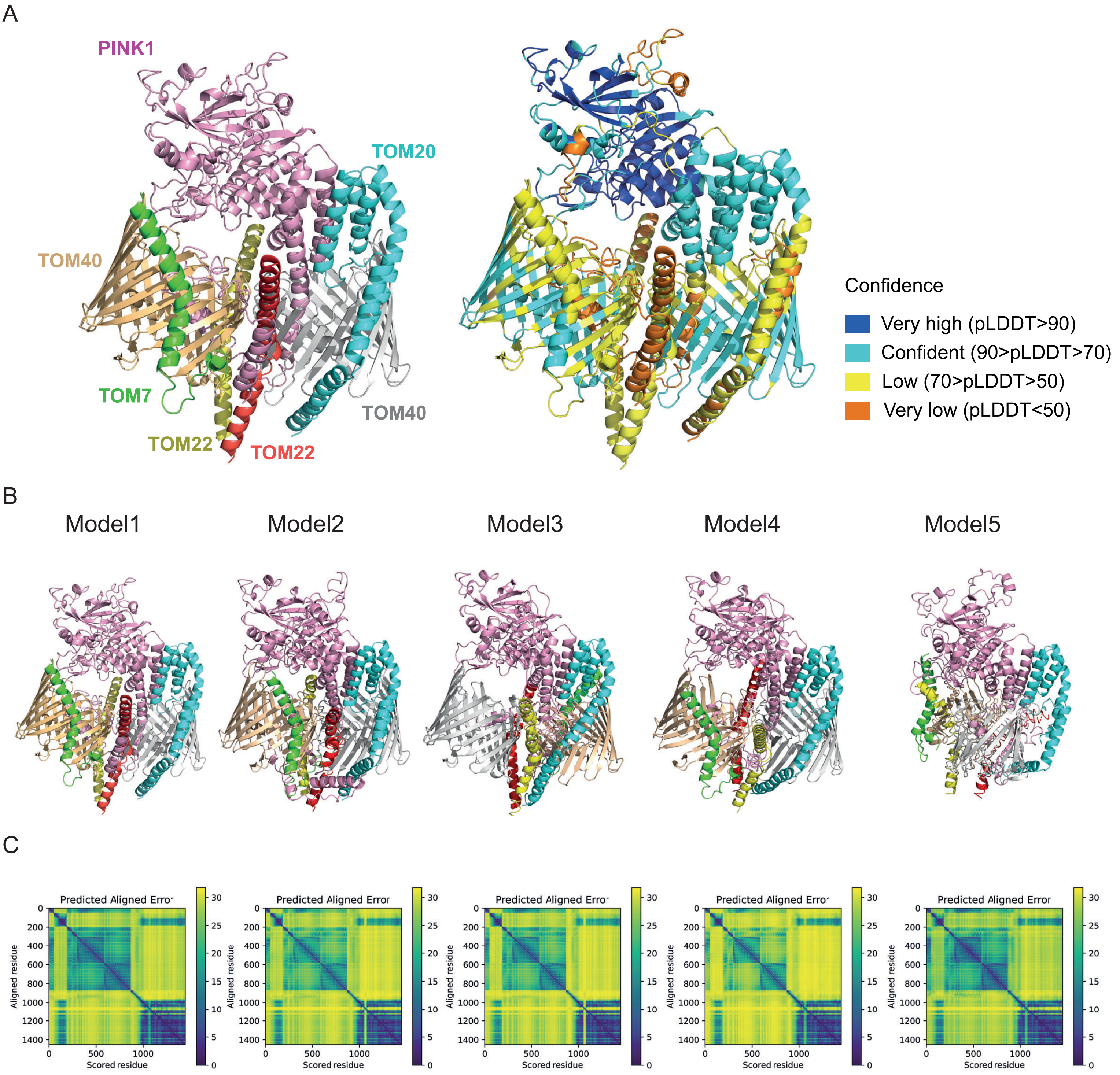

Supplementary Figure 7

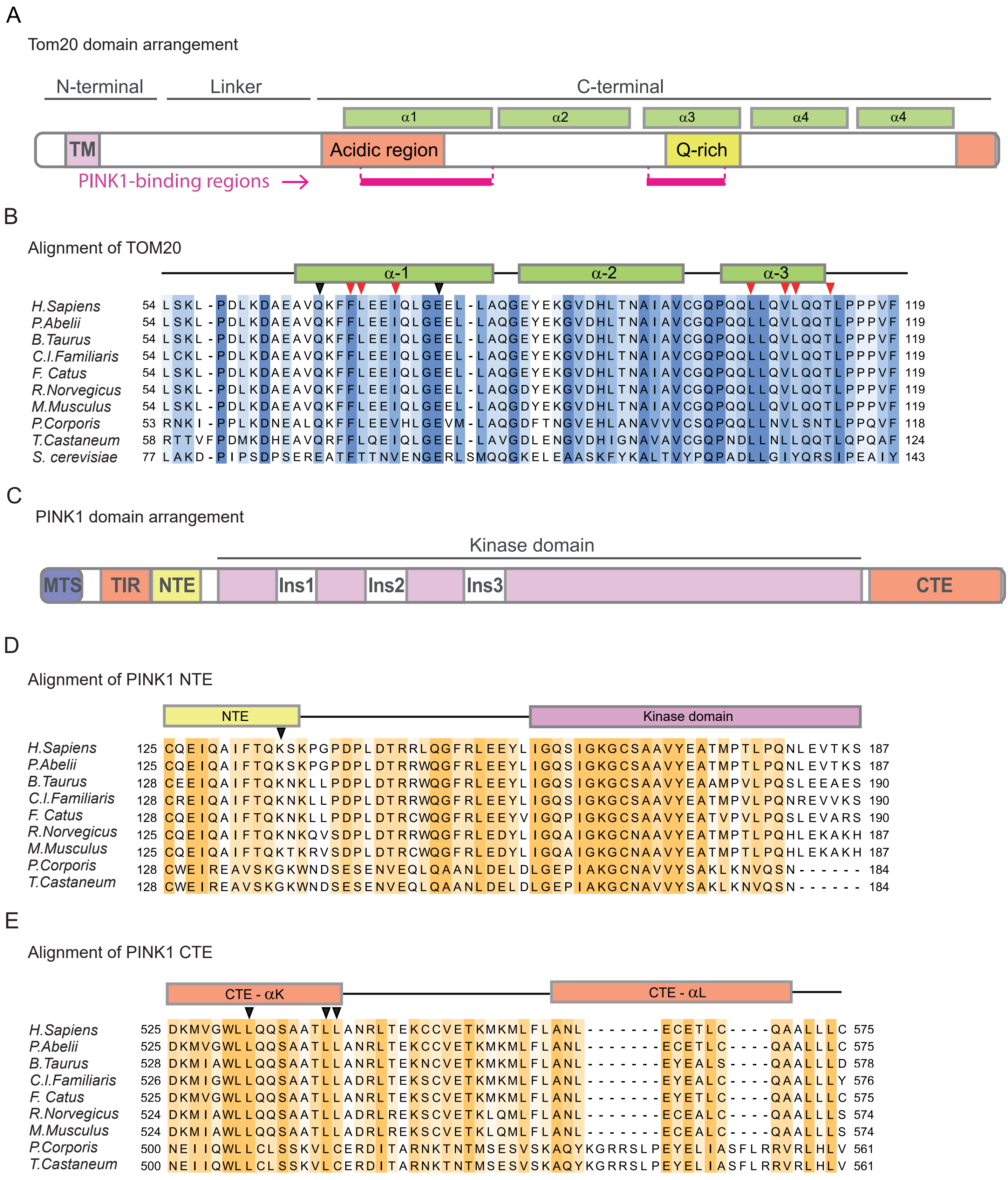

Supplementary Figure 8

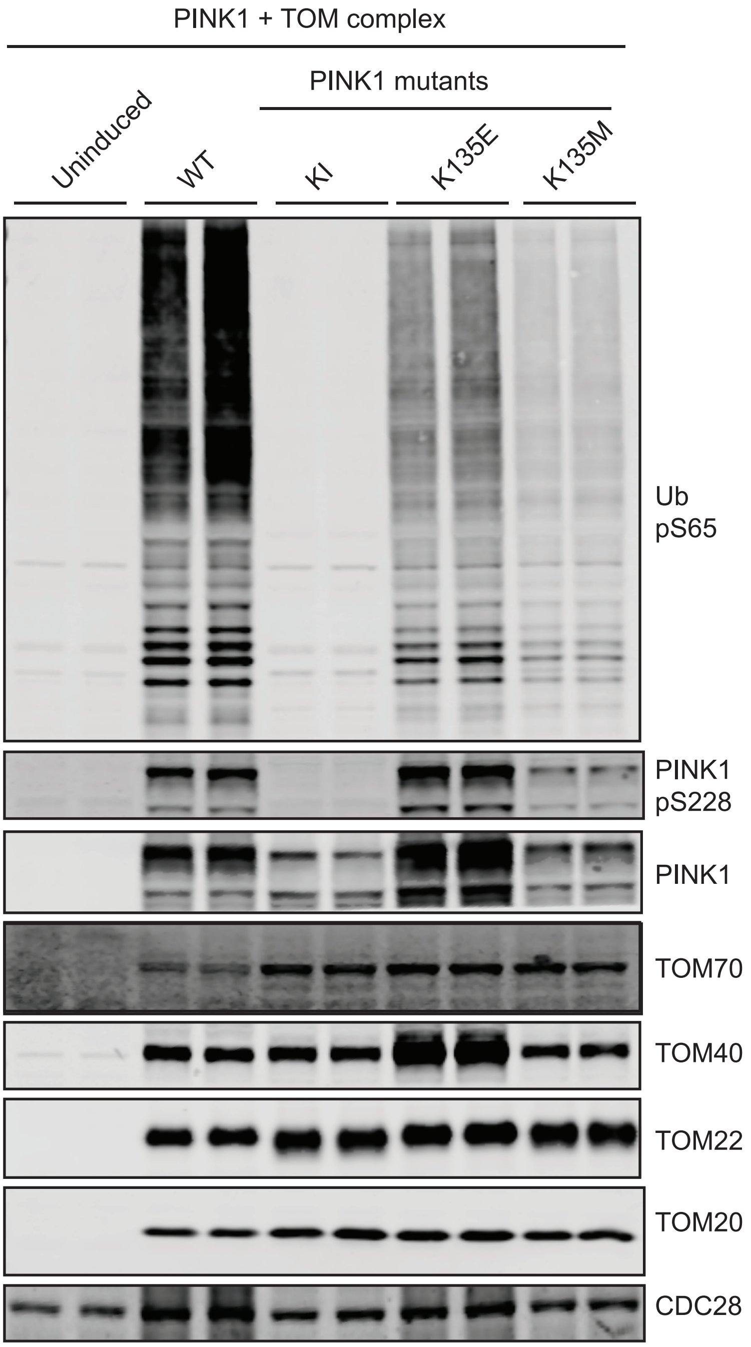

Supplementary Figure 9

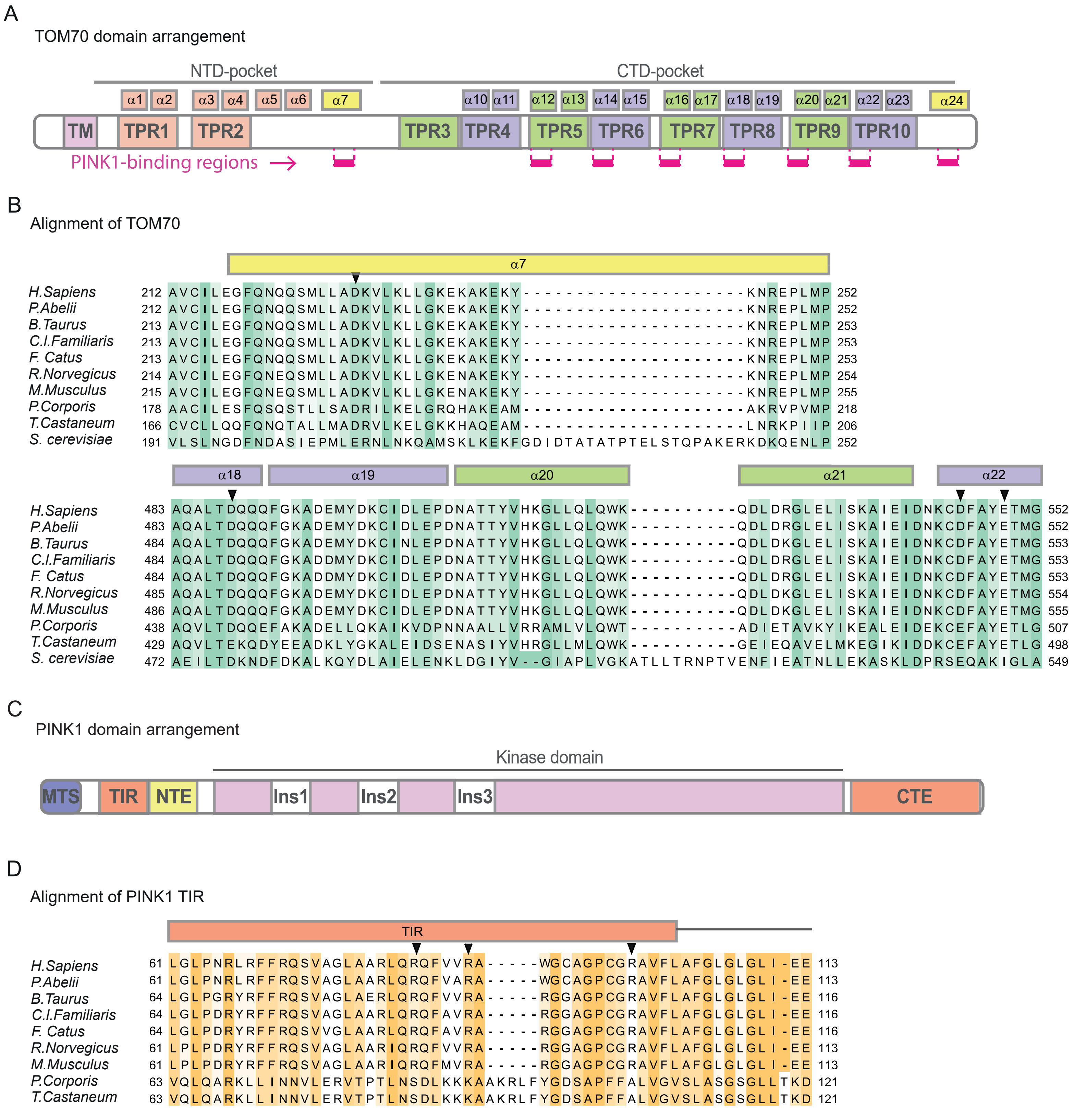

Supplementary Figure 10

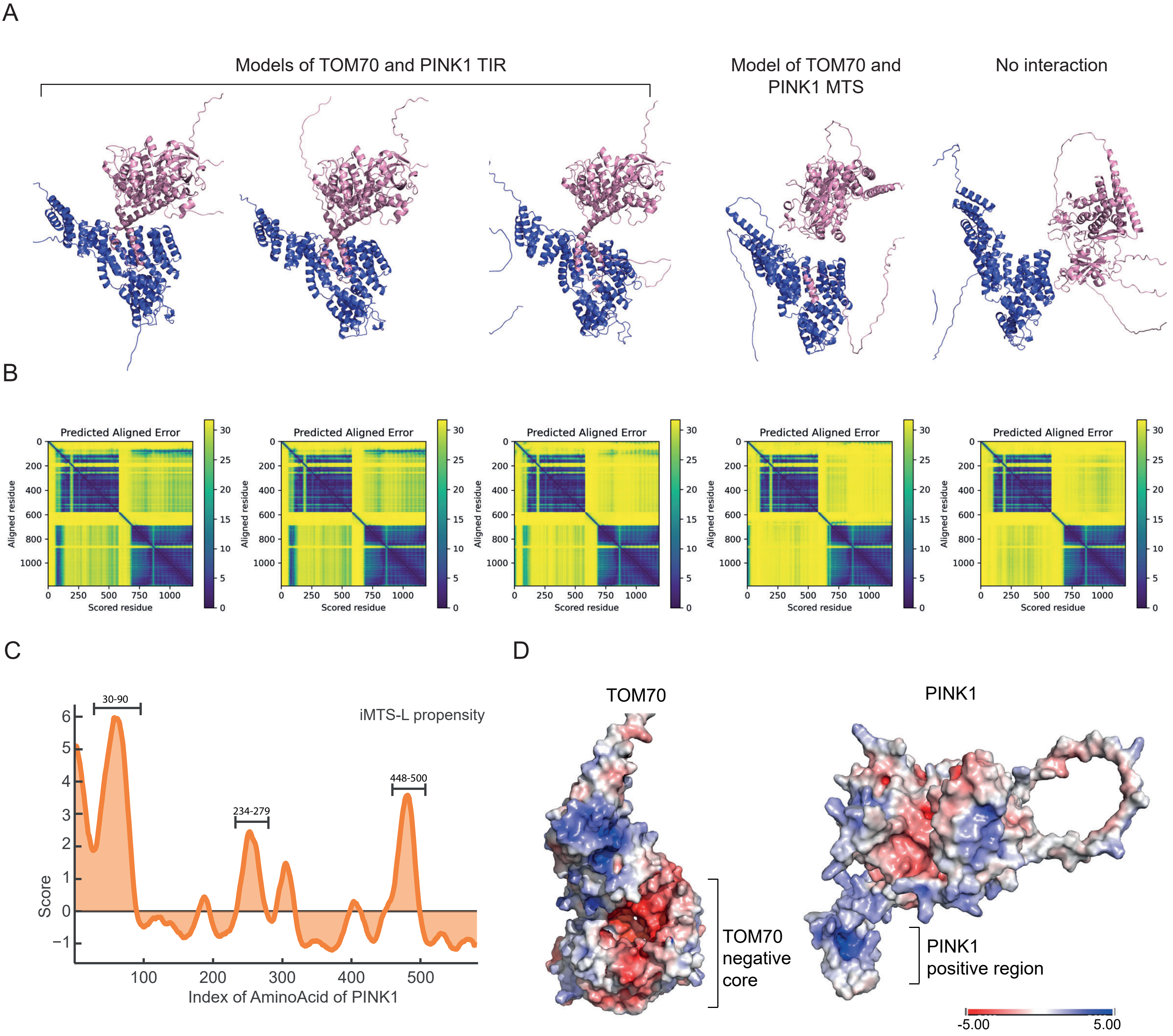

Supplementary Figure 11

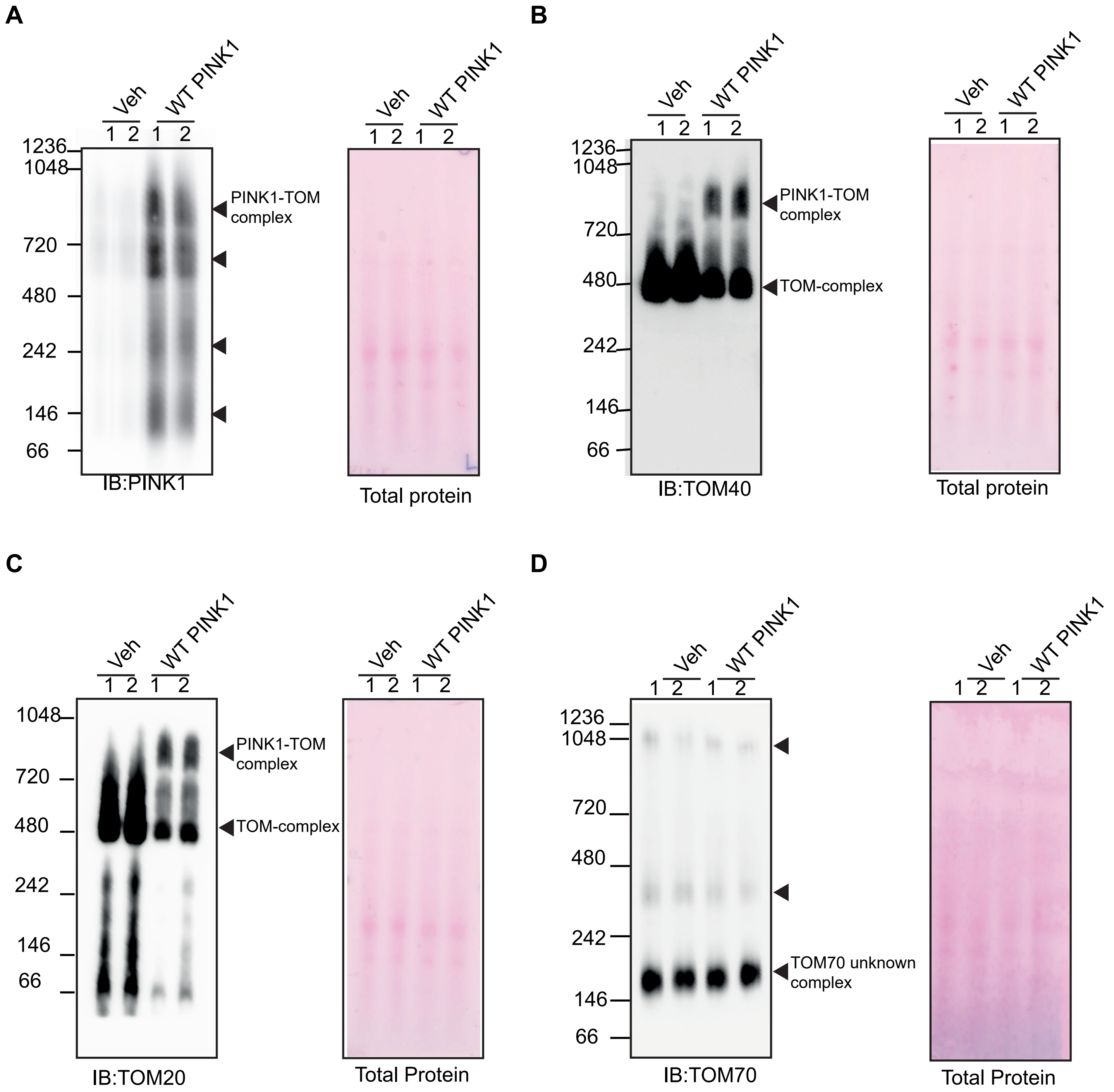

Supplementary Figure 12

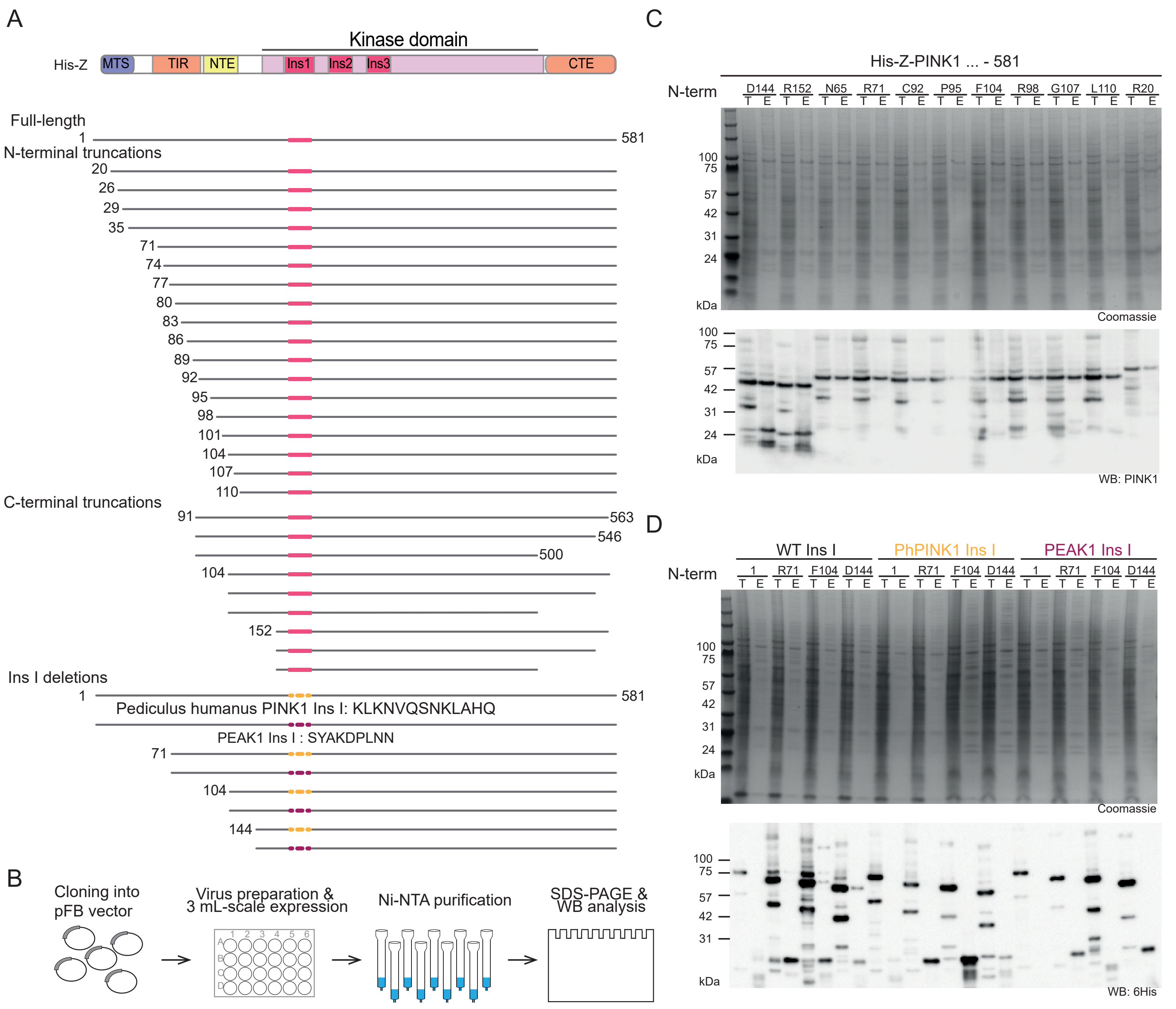

Supplementary Figure 13

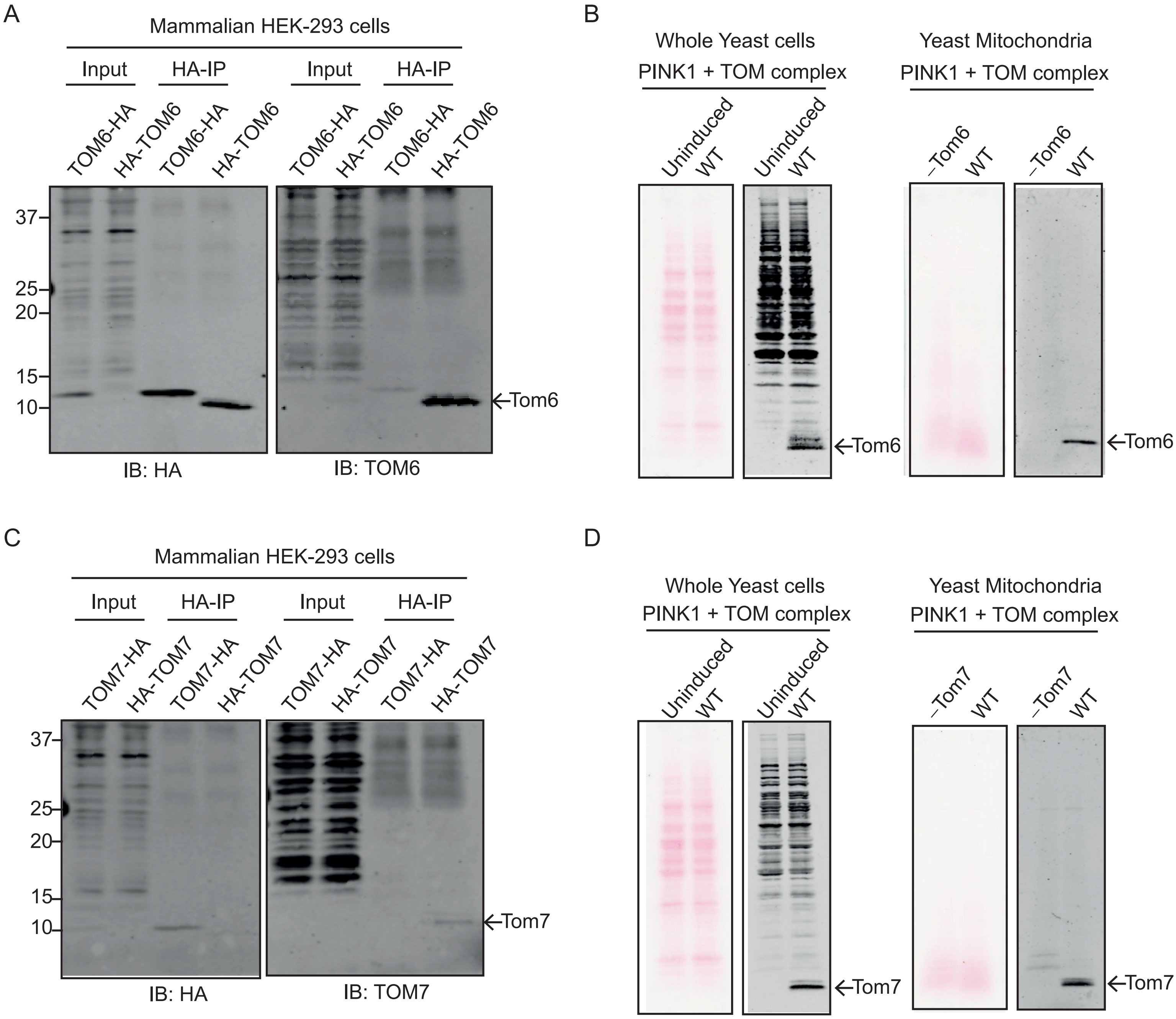
